# Environmental Novelty Modulates Rapid Cortical Plasticity During Navigation

**DOI:** 10.1101/2025.10.21.683723

**Authors:** Alexander Attinger, Antonia Drinnenberg, Can Dong, Charu Ramakrishnan, La’Akea Siverts, Tanya L Daigle, Bosiljka Tasic, Hongkui Zeng, Sean Quirin, Karl Deisseroth, Lisa M Giocomo

**Author notes:** Co-corresponding authors: Alexander Attinger; Lisa M Giocomo. These authors contributed equally (Alexander Attinger, Antonia Drinnenberg).

## Abstract

In novel environments, animals quickly learn to navigate, and position-correlated spatial representations rapidly emerge in both the retrosplenial cortex (RSC) and primary visual cortex (V1). However, the role of plasticity in building these spatial representations, and how experience modulates this process, are not well understood. Here, we investigated the plasticity of spatial representations with real-time, cellular-resolution read and write control of neural activity using two-photon calcium imaging combined with holographic optogenetic stimulation in mice navigating virtual reality environments. Targeted stimulation of individual layer 2/3 neurons rapidly biased neural activity towards stimulation-paired locations in novel, but not familiar, environments. In contrast, RSC layer 5 neurons exhibited stimulation-induced plasticity regardless of environmental familiarity. These findings reveal a layer-specific, experience-dependent modulation of plasticity and offer a framework for how neocortical spatial representations strike a balance between stability of familiar environments with flexibility for continuous updates of relevant context information.

**Highlights:**

- All optical read-write approach reveals experience-dependent plasticity in cortex
- Stimulating layer 2/3 neurons biases activity in novel but not familiar environments
- Layer 5 neurons exhibit plasticity regardless of environmental familiarity

## INTRODUCTION

Navigating animals must continuously incorporate new information about the environment, while preserving existing internal representations of familiar contexts. This balance between updating and maintaining information enables animals to learn from experience, adjust to sensory changes and optimize decision making – key adaptive behaviors essential for survival. However, the precise mechanisms that regulate neural plasticity to incorporate new information without disrupting established representations, and the timescales over which these processes can occur, remain incompletely understood. Moreover, although cellular plasticity dynamics have been extensively studied in various brain regions, our understanding of such dynamics during behavior remains limited, particularly in neocortical areas. Here, we address these gaps by investigating retrosplenial cortex (RSC), a region central to navigation in both humans and rodents (Sutherland, Whishaw, and Kolb 1988; Takahashi et al. 1997; Harker and Whishaw 2002; Vann and Aggleton 2002) and primary visual cortex (V1), a direct input to RSC (Wang and Burkhalter 2007; Oh et al. 2014; Fischer et al. 2020).

The RSC serves as a hub for integrating visual and spatial information (Campbell et al. 2021; Mao et al. 2020; Fischer et al. 2020), with strong connectivity not only to visual cortical areas but also the hippocampal formation (Sugar et al. 2011; Qiu et al. 2024; Lin et al. 2024) and medial entorhinal cortex (Sürmeli et al. 2015; Simonsen, Czajkowski, and Witter 2022; Dubanet and Higley 2024), two brain regions deeply involved in navigation (Parron and Save 2004; Robinson et al. 2020). Similar to the hippocampus (O’Keefe and Dostrovsky 1971), RSC neurons can exhibit position-correlated activity patterns (Alexander and Nitz 2015; Mao et al. 2017; Campbell et al. 2021) that stabilize with experience (Mao et al. 2018; Fischer et al. 2020; Miller, Mau, and Smith 2019), and encode spatial context information (Smith et al. 2004; Cowansage et al. 2014). Like RSC, V1 neural activity can also be spatially modulated, and exhibits coordinated activity with the hippocampus that, in turn, can shape some forms of experience dependent plasticity (Ji and Wilson 2007; Haggerty and Ji 2015; Fiser et al. 2016; Pakan et al. 2018; Saleem et al. 2018; Finnie, Komorowski, and Bear 2021).

In the hippocampal region CA1, the formation of stable place-specific firing (i.e. place cells) is thought to be driven in part by behavioral timescale synaptic plasticity (BTSP) (Bittner et al. 2017). This mechanism enables synaptic weight changes leading to place field formation on the timescale of seconds to minutes. Whether similarly rapid plasticity mechanisms operate in neocortical regions like RSC however, and the degree to which experience modulates such plasticity, remain unknown. To investigate the processes and timescales underlying position encoding in RSC and V1, we combined two-photon calcium imaging with millisecond-precision holographic optogenetic stimulation of groups of neurons at single-cell resolution in mice navigating novel and familiar virtual reality (VR) environments. We selectively activated groups of single neurons at pre-defined spatial locations, a strategy known to induce place fields in hippocampal region CA1 (Rolotti et al. 2022; O’Hare et al. 2022; Geiller et al. 2022; Fan et al. 2023). This allowed us to investigate if a mechanism based on rapid plasticity contributes to the formation of spatial representations in RSC and V1 in an experience-dependent manner.

Our findings reveal that repeated stimulation of cortical neurons at specific VR locations rapidly modifies their activity within a single session, in an experience-dependent manner. Specifically, in novel environments, stimulation rapidly induced a long-term increase in activity at the stimulation-paired position in a subset of targeted neurons, whereas the same manipulation had no long-term effect in familiar environments. This experience-dependent plasticity was present in the cortical layer 2/3 (L2/3) of both RSC and V1, thus generalizable across cortical areas. In contrast to L2/3, layer 5 (L5) RSC neurons exhibited stimulation-induced plasticity independent of prior experience, as stimulation induced long-term increases in activity at stimulation-paired positions even in familiar environments. These results provide evidence for rapid plasticity mechanisms in both RSC and V1 and reveal a layer-specific hierarchy in how experience determines the temporal window of plasticity in the cortex, allowing neocortical circuits to strike a balance between flexibility and stability.

## RESULTS

### Probing fast timescale plasticity in cortical neurons during spatial navigation

We investigated the plasticity of spatial representations in RSC and V1 by recording and manipulating neuronal activity across three consecutive days in transgenic mice that expressed the calcium indicator GCaMP6m together with the excitatory opsin ChRmine in excitatory cortical neurons (*Ai228;CamK2a-CreERT2* (Drinnenberg et al. 2024), Methods). Head-fixed mice navigated a VR track with patterned elements providing optic flow, fixed visual cues and two transient landmarks providing position information (LM1, LM2, Fig. 1A, B). To precisely time the onset of stimulation with the visibility of LM1 and LM2, they appeared next to the mouse for 1 second at predefined locations along the VR track and moved in synchrony with the optic flow pattern. The experiment consisted of three sessions over three days: a baseline session on Day 1 (10 trials), a stimulation session on Day 2 (30 stimulation trials, followed by ≥ 10 non-stimulation trials), and a post-stimulation session on Day 3 (10 trials), allowing analysis of rapid and long-term activity changes. To examine the influence of experience, we also varied the prior exposure to visual VR: Naïve mice experienced visual VR for the first time on Day 1, while familiar mice had prior VR experience (7 or more sessions, Methods). These experience states were reflected in the behavior over the three session days: Naïve mice learned the location of the reward across the three days, as indicated by behavioral changes in licking and running speed (Fig. S1A, B), whereas behavior in familiar mice was stable (Fig. S1C, D).

**Figure 1:**
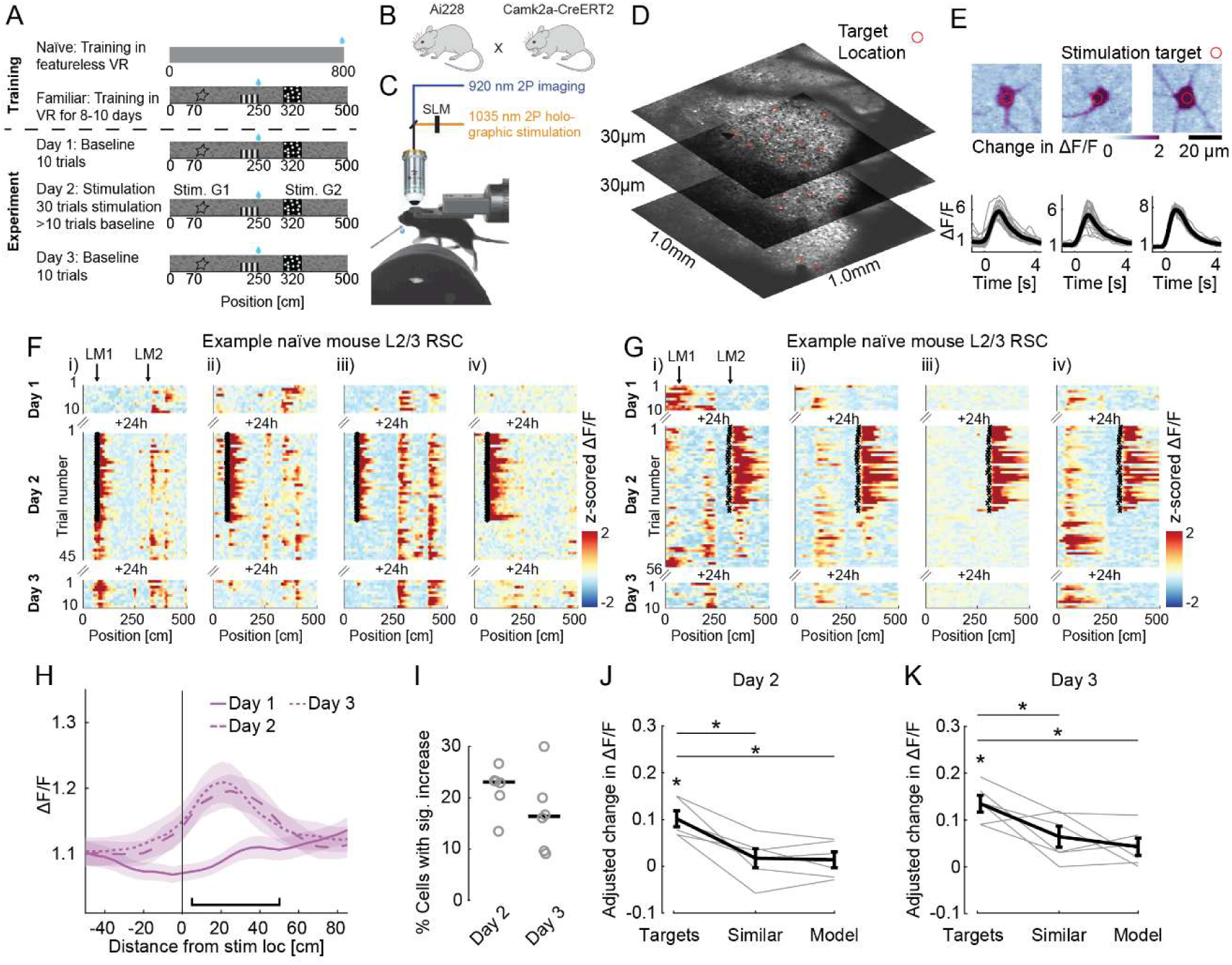
Rapid and lasting spatial activity bias induced by targeted stimulation of L2/3 RSC neurons. A. Schematic of the training and experiment protocol. Naïve mice were trained in a featureless VR, with progressively increasing distances between rewards. Familiar mice experienced the VR for 7-10 sessions before the experiment. The experiment consisted of three sessions. On Day 1 (Baseline), mice were exposed to VR for 10 trials. On Day 2, holographic stimulation was applied to two groups of neurons (G1 and G2). This stimulation (Stim.) was paired with the appearance of visual landmarks and lasted 30 trials. Mice then ran an additional set of trials immediately after with no stimulation (n ≥ 10 post-stimulation trials). On Day 3, mice ran 10 trials with no holographic activation B. We used a transgenic approach (*Ai228;CamK2a-CreERT2*) to express GCaMP6m and soma-enriched ChRmine-oScarlet in excitatory neurons throughout cortex. Cre recombination was induced by administration of tamoxifen prior to the surgery (Methods). C. Schematic of the setup for simultaneous 2P imaging and 2P holographic stimulation in behaving animals. D. Example of a multiplane imaging field of view showing L2/3 in agranular RSC of an example *Ai228;CamK2a-CreERT2* mouse, with red circles indicating target locations. E. Stimulation response of three example stimulation targets. Top: ΔF/F images of the stimulation-evoked activity, centered around 3 stimulation targets. Red circle indicates outer limit of targeting spiral (diameter: 7 µm). Scale bar: 20 µm. Bottom: Single trial (gray lines) and trial-averaged (black line) stimulation-evoked activity of the same neurons (n = 30 stimulation trials). F. Spatially binned activity of four co-recorded Target neurons from group 1 (G1) of a naïve mouse with stimulation paired only to landmark 1 (LM1) on Day 2. Arrows on top indicate the position of the stimulation-paired landmarks. Small black x’s mark the onset of stimulation. Note that activity around LM1 is significantly increased in Target neurons on the left (i and ii), but unchanged in iii and iv (i, ii: p < 0.025, iii, iv: p > 0.05 one-tailed Wilcoxon signed-rank test, comparing LM1 surrounding activity on 10 post stimulation trials to activity on Day 1). G. Same as in (F), but for four target neurons from group 2 (G2) from a different naïve mouse, with stimulation paired to landmark 2 (LM2). Activity around LM2 is significantly increased in Target neurons on the left (i and ii), but unchanged in iii and iv (i, ii: p < 0.025, iii, iv: p > 0.05 one-tailed Wilcoxon signed-rank test, comparing LM2 surrounding activity on 10 post stimulation trials to activity on Day 1). H. Mean spatially binned activity around the stimulation-paired landmark of all Target neurons in naïve mice. Data from LM1 and LM2 target neurons are combined (n = 260 Target neurons, 6 mice, median = 46, range = 30-56 Target neurons per mouse). Day 1 (solid line), Day 2 after stimulation trials ended (long dash) and Day 3 (short dash). Shading indicates SEM. Black horizontal line: ‘landmark-surrounding’ region used for quantification in subsequent figures (5 - 50 cm post onset position). I. The fraction of Target neurons per mouse (n = 6 mice) with a significant increase in landmark-surrounding activity. Left: Comparing Day 2 activity to Day 1 activity. Right: Comparing Day 3 activity to Day 1 activity. Grey circles: average fraction per mouse, black line; median. J. Adjusted change in landmark-surrounding activity per mouse between Day 1 and Day 2 for Target neurons (left), compared to co-recorded neurons with similar activity on Day 1 (middle, ‘Similar’), as well as Model neurons trained to predict the activity of Target neurons on Day 1 (right, ‘Model’) (Methods). Adjusted change was computed for each animal by subtracting the change in landmark-surrounding activity in the left-out population from the observed changes in Target, Similar, and Model neurons. Change in activity in Target neurons is significantly larger than Similar neurons and Model neurons. (Targets vs. Similar: p ≤ 0.03, Targets vs. Model: p ≤ 0.05, Targets vs. 0: p ≤ 0.02, one-sided Wilcoxon signed-rank test, n = 6 mice). Black line indicates mean ± SEM, gray lines indicate mean change per mouse. K. Same as in (J), but comparing activity between Day 1 and Day 3 (Targets vs. Similar: p ≤ 0.05, Targets vs. Model: p ≤ 0.03, Targets vs. 0: p ≤ 0.02, one-sided Wilcoxon signed-rank test, n = 6 mice). *p ≤ 0.05, **p ≤ 0.01

We used a custom-built spatial light modulator (SLM) integrated into a two-photon microscope to activate groups of targeted neurons with high spatial and temporal resolution spread throughout the imaging volume (Fig. 1C-E Fig. S1E-H). In each mouse, we selected two groups of neurons based on their activity patterns during the Day 1 baseline session (group 1 and group 2). Group 1 target neurons were required to exhibit low activity surrounding LM1 during the baseline session (LM2 for group 2 neurons, respectively). During the stimulation session on Day 2, we paired the stimulation of group 1 and group 2 neurons with the appearance of LM1 or LM2, respectively. Neurons were stimulated for 30 trials (1 second stimulation at 31 Hz) at the beginning of the session either consecutively, or with three no-stimulation trials interleaved (trials 8, 18 and 28 on Day 2). Stimulation trials were followed by at least 10 non-stimulation trials to assess rapid stimulation-induced changes, and activity was recorded from the same neurons on Day 3 to assess longer lasting changes.

### Stimulation induces lasting changes in RSC neural activity in naïve mice

We first investigated if location-paired activation of targeted L2/3 neurons in agranular RSC (Paxinos and Franklin 2001) could bias spatial selectivity in naïve mice (i.e. mice with no exposure to visual VR environments before the Day 1 baseline session). Putative target neurons were defined based on the activity during the baseline session, as described above, and stimulated on Day 2. We confirmed that stimulation strongly activated selected target neurons (Fig. S1G, H). On average, 79% of selected neurons were reliably and strongly activated during the stimulation session and tracked across the three imaging sessions (‘Target neurons’ median = 23, range = 14 - 30 Target neurons per group, 260 of 329 selected neurons). The strong activation of Target neurons was accompanied by a small decrease in activity in non-stimulated ‘left-out’ neurons (distance to closest stimulation target ≥ 15 μm, Fig. S1I).

We then analyzed the post-stimulation trials on Day 2 to evaluate lasting effects of stimulation on the Target neuron population. Compared to Day 1, average activity in Target neurons increased around the stimulation-paired location (Fig. 1F-H, Fig. S1J). This bias in activity was significant in 55 out of 260 Target neurons (21%, range = 14-29%, median = 23% per mouse, n = 6 mice; Fig. 1I), and was already observable during the interleaved trials (Fig. S1K, L), indicating that location-paired activation induced significant and rapid changes in the Target neurons.

To control for both the effect of the selection criteria, and normally occurring drift in spatially correlated activity, we compared the changes in the activity of Target neurons to changes in activity-matched co-recorded neurons (’Similar’) and models trained to predict the activity of Target neurons (’Model’). The change in activity for Target neurons in post-stimulation trials was larger than the change in activity for both the Similar and Model groups (Fig. 1J, Fig. S1M), indicating that the observed effects are unlikely to be caused by selection bias or normally occurring drift alone. We then asked whether the induced activity fields in post-stimulation trials in Target neurons were similar to activity fields observed in left-out neurons. We found that the peak amplitude and spatial extent of induced fields in Target neurons were similar to activity fields observed in left-out neurons (Fig. S1N, O). Together, these data demonstrate that neural activity can be biased towards stimulation-paired locations in a subset of L2/3 RSC neurons within a single session.

To assess persistent changes in activity across days, we analyzed the activity of Target neurons on Day 3, 24 hours after stimulation trials on Day 2. Remarkably, activity remained increased around the stimulation-paired landmarks (Fig. 1H, K, Fig. S2A, B) and the fraction of neurons with significantly increased activity compared to baseline was comparable between Day 2 and 3 (median Day 2 = 23 %, median Day 3 = 16%, Wilcoxon signed-rank, p > 0.05, n = 6 mice). Moreover, the change in landmark-surrounding activity of Target neurons was highly correlated across days, indicating that Target neurons with an increase in activity on Day 2 retained this increased activity on Day 3 (correlation = 0.73, p < 0.001, n = 260 Target neurons, Fig. S2C). Interestingly, we did not observe a backward shifting of the induced landmark-surrounding activity on Day 2 or Day 3 (Fig. S2D), a feature that has been associated with behavioral timescale plasticity in CA1 (Bittner et al. 2017; Rolotti et al. 2022).

While activity remained increased in Target neurons across days, not all Target neurons exhibited significant bias in activity. We therefore analyzed if the stimulation-induced effect in individual neurons could be predicted based on their functional properties. Neither running modulation nor spatial stability during the baseline session was predictive of the stimulation-induced bias in Target neurons (multiple linear regression: R^2^: 0.003, F(2,257) = 0.332, p = 0.7). We also investigated if the evoked effects depended on varying stimulation parameters across Target neurons, as previous work has shown that artificial induction of place fields depends on the number of simultaneously targeted CA1 neurons (Rolotti et al. 2022). However, we found no relationship between the change in activity in Target neurons and the distance to the next closest target (R = 0.002, F(2,257) = 0.3, p = 0.7, multiple linear regression), or the total number of simultaneously activated targets (Kruskal-Wallis H(7) = 13.0, p = 0.07). These results therefore suggest inherent heterogeneity in the plasticity of RSC L2/3 neurons in novel environments.

Finally, we examined if activity in Target neurons significantly changed outside of the stimulation paired landmark. We analyzed their activity at all positions except the region surrounding the stimulation-paired landmark by excluding -10 cm to 60 cm around LM1 (LM2) for group 1 (group 2) Targets. In contrast to the increase in activity around the stimulation-paired landmark, activity outside the stimulation-paired landmark decreased in post-stimulation trials (Fig. S2E). Moreover, there was no correlation between this decrease and the increase in activity around the stimulation-paired landmark on Day 2 (rho: 0.07, p ≥ 0.25, n = 260 Target neurons), or on Day 3 (rho: 0.07, p ≥ 0.28, n = 260 Target neurons). Finally, in Target neurons, the decrease of activity outside the stimulation-paired landmark on Day 3 was comparable to that observed in Similar neurons (Fig. S2F). As this decrease in activity appeared unrelated to the lasting increase in activity at the stimulation-paired landmark, it is unlikely to reflect a redistribution of neural activity towards the stimulation-paired landmark. Rather, it could reflect short-term effects on intrinsic excitability in the stimulated neurons. Collectively, these findings demonstrate that targeted holographic stimulation can induce rapid and persistent activity biases in L2/3 RSC neurons during initial VR environment exploration.

### Stimulation-induced plasticity in L2/3 of RSC is experience dependent

We then asked whether RSC representations become less susceptible to optogenetic perturbations as environments become more familiar, given that spatial representations in RSC stabilize with experience (Fischer et al. 2020; Mao et al. 2018). To address this question, we repeated the stimulation experiments in mice that were trained in the VR environment for at least 7 days prior to the Day 2 stimulation experiment. In contrast to the effects in naïve mice, activity surrounding the stimulation-paired landmark was unchanged in Target neurons of familiar mice in post-stimulation trials on Day 2, and increased only slightly on Day 3 (Fig. 2A-D, Fig. S2G-I, 208 Target neurons, median = 22, range = 11-32 Target neurons per landmark, 29-58 Target neurons per mouse, n = 5 mice). This was despite the strong activity modulation during stimulation trials that was comparable to effects seen in naïve mice (Fig. S2J-L). In contrast to L2/3 of RSC, where stimulation induced plasticity appears to be modulated by experience, we were able to induce place fields in CA1 neurons in mice that were familiar with the environment (Fig. S2M, N).

**Figure 2:**
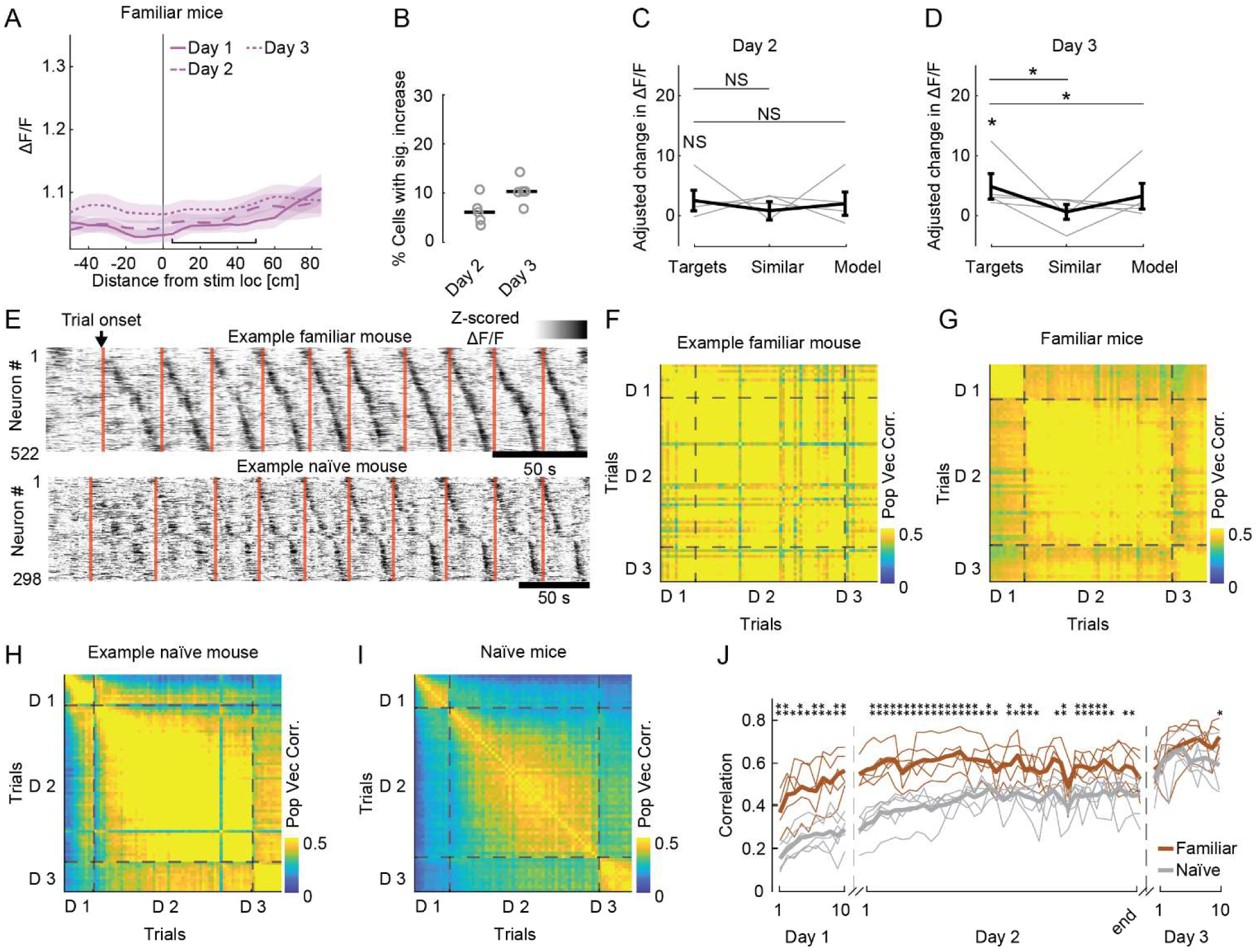
Familiarity modulates plasticity in RSC. A. Mean spatially binned activity around the stimulation-paired landmark of all Target neurons in familiar mice. Data from LM1 and LM2 Target neurons are combined (n = 208 neurons, 5 mice, range = 28 to 58 Target neurons per mouse). Day 1 (solid line), Day 2 after stimulation trials ended (long dash) and Day 3 (short dash). Shading indicates SEM. Black horizontal line: ‘landmark-surrounding’ region used for quantification. B. The fraction of Target neurons per mouse (n = 5 mice) with a significant increase in landmark-surrounding activity on Day 2 and Day 3, relative to activity on Day 1 (as in Fig. 1I). Grey circles: average fraction per mouse; black line: median. C. No significant change in activity around the stimulation-paired landmark between Day 1 and Day 2 for Target neurons (left), compared to Similar and Model neurons (as in Fig. 1J, Targets vs. Similar: p > 0.05, Targets vs. Model: p > 0.05, Targets vs. 0 p > 0.05, Wilcoxon signed-rank test, n = 5 mice). Black line indicates mean ± SEM, gray lines indicate mean change per mouse. D. Same as (C), but for the difference in activity between Day 1 and Day 3. Increase in landmark-surrounding activity in Target neurons is larger than in Similar and Model neurons (Targets vs. Similar: p ≤ 0.03, Targets vs. Model: p ≤ 0.03, Targets vs 0: p ≤ 0.03, one-sided Wilcoxon signed-rank test, n = 5 mice). E. Top: Activity (z-scored ΔF/F) of position-stable RSC L2/3 neurons (n = 522 neurons) from the left-out population of an example familiar mouse during baseline trials on Day 1. Neurons are ordered by the preferred position on Day 2. Red lines indicate trial onset. Bottom: Same as top, but for the position-stable neurons of an example naïve mouse exploring visual VR for the first time (n = 298 neurons). F. Spatial similarity of neural activity across trials from an example familiar mouse (same neurons as in (E, top)). Heatmap shows population vector correlation of position-stable left-out neurons (n = 522 neurons), comparing activity between all trials within and across three experimental days (D1, D2, D3). Dashed lines indicate day boundaries. G. Average spatial similarity of neural activity across trials in all familiar mice (n = 5 mice, median = 466, range = 335-609 neurons). H. Same as in (F), but for the spatial similarity of neural activity across trials from an example naïve mouse (same neurons as in (E, bottom), n = 298 neurons). I. Average spatial similarity matrix for all naïve mice (n = 6 mice, median = 368, range = 157-533 position-stable left-out neurons). J. Correlation of spatial activity patterns of left-out neurons between Days 1 and 2 and Day 3 in familiar (brown) and naïve (gray) mice. Higher correlation indicates greater similarity to Day 3. Thick lines: mean; thin lines: individual mice. *p < 0.05, **p < 0.01 (Mann-Whitney U test, n = 6 naïve mice, 5 familiar mice). Day 2 data is interpolated to accommodate varying trial numbers (Methods).

While stimulation did not appear to rapidly change activity surrounding the stimulation-paired landmark, Target neuron activity decreased outside the stimulation-paired landmark on Day 2, and recovered on Day 3 (Fig. S2O, P). Notably, this temporary decrease on Day 2 was similar to the reduction we observed in naïve mice, further indicating that stimulation can also lead to a temporary decrease in activity.

Finally, we analyzed the spatially correlated activity in left-out neurons in both naïve and familiar mice. In line with previous work, we observed that the neural representation of the familiar VR environment by left-out neurons remained highly stable across the three days of the experiment (Fig. 2E-G). In contrast, in naïve mice, spatial representations in left-out neurons were initially unstable, stabilized across days and reached similar stability levels as the familiar mice only on Day 3 of imaging (Fig. 2E, H - J). These results further highlight the rapid plasticity of spatial representations in RSC during the initial exposure to a novel environment and demonstrate the subsequent stability of these representations once the environment becomes familiar.

### Environmental novelty modulates L2/3 plasticity across cortical areas

RSC is a cortical hub for integrating visual and spatial information, raising the question of whether the rapid and experience-dependent plasticity we observed is a specific property of RSC, or is generalizable across cortical areas. To address this question and directly compare the plasticity of spatially modulated activity across cortical areas, we next conducted identical experiments in L2/3 of V1. As in RSC, pairing visual landmarks with stimulation on Day 2 in naïve mice led to an increase in landmark-surrounding activity in post-stimulation trials in V1 Target neurons (Fig. 3A-E, Fig. S3A, B; 338 selected neurons, 250 reliably activated and tracked Target neurons, range 4-25 Target neurons per landmark, range 15-43 Target neurons per mouse, 8 naïve mice). The induced bias in landmark-surrounding activity was already observable during the interleaved trials (Fig. S3C, D), persisted across days (Fig. 3F, S3E - G, correlation between Day 2 change and Day 3 change: rho = 0.47, p<0.001, n = 250 neurons), and was similar to activity in left-out neurons (Fig. S3H, I). This indicates that stimulation-induced changes occurred on a similarly fast timescale compared to RSC.

**Figure 3:**
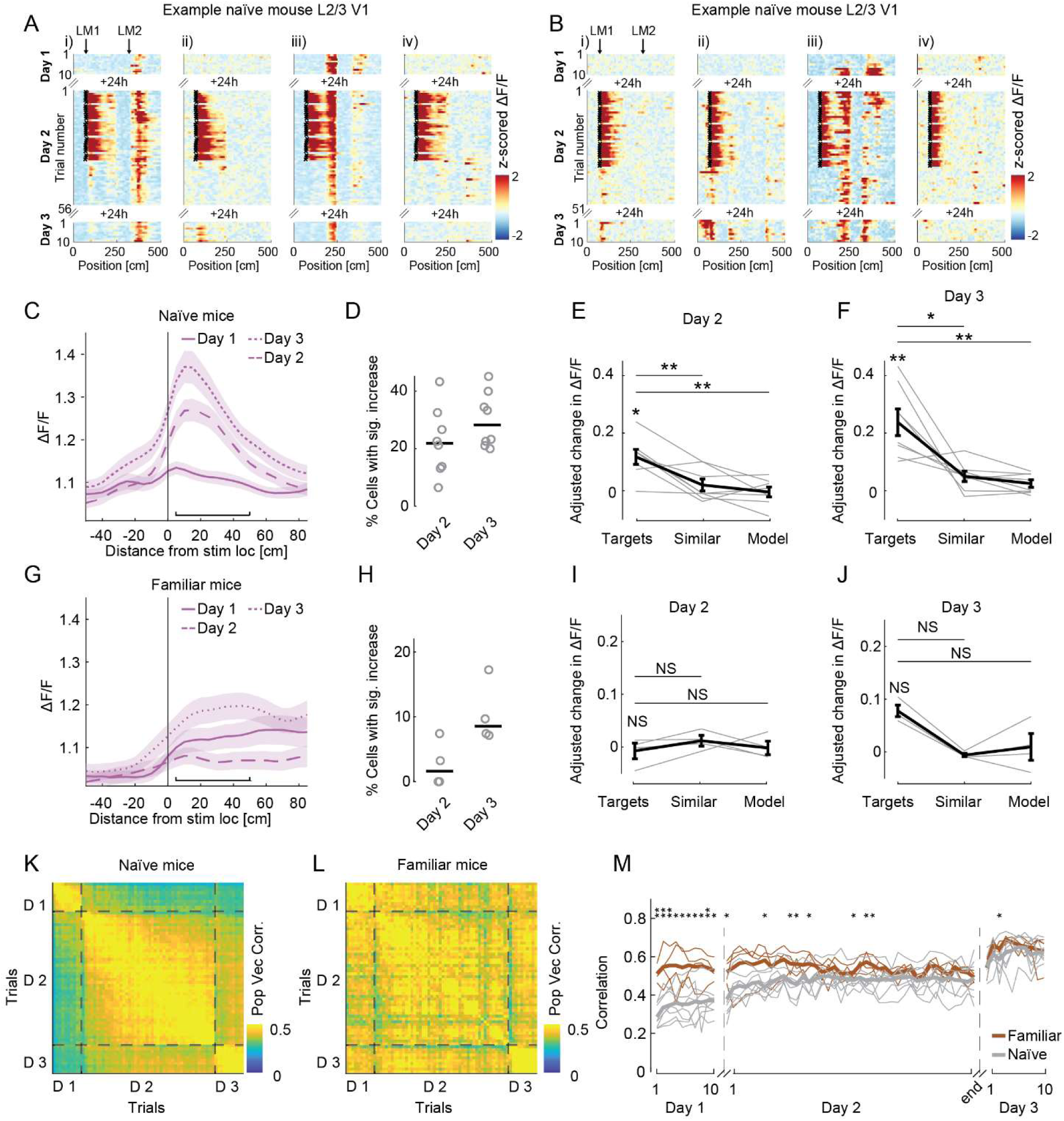
Novelty gates plasticity in L2/3 neurons of V1. A. Spatially binned activity of four co-recorded Target neurons from V1 of a naïve mouse. Target neurons are from group 1, with stimulation paired only to landmark 1 (LM1) on Day 2. Activity around LM1 is significantly increased in Target neurons on the left (i and ii), but unchanged in iii and iv (i, ii: p < 0.025; iii, iv: p > 0.05 one-tailed Wilcoxon signed-rank test, comparing LM1 surrounding activity on 10 post-stimulation trials to activity on Day 1). Arrows on top indicate the position of stimulation-paired landmarks. Small black x’s mark the onset of stimulation. B. Same as in (A), but for four Target neurons from a different mouse. Activity around LM1 is significantly increased in Target neurons on the left (i and ii), but unchanged in iii and iv (i, ii: p < 0.025; iii, iv: p > 0.05 one-tailed Wilcoxon signed-rank test, comparing LM1 surrounding activity on 10 post-stimulation trials to activity on Day 1). C. Mean spatially binned activity around the stimulation-paired landmark of all Target neurons in V1 of naïve mice. Data from LM1 and LM2 Target neurons are combined (250 neurons, 8 mice, median = 31, range = 15 to 43 Target neurons per mouse). Day 1 (solid line), Day 2 after stimulation trials ended (long dash) and Day 3 (short dash). Shading indicates SEM. Black horizontal line: ‘landmark-surrounding’ region used for quantification. D. The fraction of Target neurons per mouse (n = 8 mice) with a significant increase in landmark-surrounding activity on Day 2 and Day 3, relative to activity on Day 1 (as in Fig. 1I). E. Change in landmark-surrounding activity per mouse between Day 1 and Day 2 for Target neurons (left), compared to Similar and Model neurons. Change in activity in Target neurons is significantly larger than in Similar and Model neurons. (Targets vs. Similar: p ≤ 0.02, Targets vs. Model: p ≤ 0.02, Targets vs. 0: p ≤ 0.02, one-sided Wilcoxon signed-rank test, n = 8 mice). Black line indicates mean ± SEM, gray lines indicate mean change per mouse. F. Same as in (E), but comparing activity between Day 1 and Day 3 (Targets vs. Similar: p ≤ 0.02, Targets vs. Model: p ≤ 0.01, Targets vs 0: p ≤ 0.01, one-sided Wilcoxon signed-rank test, n = 8 mice). G. Same as in (C), but for Target neurons in V1 of familiar mice (n = 115 Target neurons, 4 mice). H. Same as in (D), but for the fraction of Target neurons with significant change in familiar mice (n = 4 mice). I. Same as in (E), but for adjusted difference in landmark-surrounding activity between Day 2 and Day 1 in familiar mice. p > 0.05 for all comparisons, one-sided Wilcoxon signed-rank test, n = 4 mice. Note that the relatively small number of animals makes the statistical test difficult to interpret. See also Fig. S4B. J. Same as in (F), but for the adjusted difference between Day 3 and Day 1. For all comparisons, p > 0.05, one-sided Wilcoxon signed-rank test, n = 4 mice. Note that the relatively small number of animals makes the statistical test difficult to interpret. See also Fig. S4G. K. Spatial similarity in non-stimulated neurons across trials in naïve mice. Heatmap shows population vector correlation of position-stable neurons, comparing activity between all trials within and across three experimental days (D1, D2, D3). Dashed lines indicate day boundaries. (n = 8 mice, median = 223, range = 114-383 position-stable neurons). L. Same as in (K), but spatial similarity matrix for all familiar mice (n = 4 mice, median = 153, range = 51-195 position-stable left-out neurons). M. Correlation of spatial activity patterns of left-out neurons between Days 1 and 2 and Day 3 in familiar (brown) and naïve (gray) mice. Higher correlation indicates greater similarity to Day 3. Thick lines: mean; thin lines: individual mice. *p < 0.05, **p < 0.01 (Mann-Whitney U test, n = 8 naïve mice, n = 4 familiar mice). Day 2 data are interpolated to accommodate varying trial numbers (Methods).

When paired with a landmark, stimulation-induced activity fields did not appear to shift backward (Fig. S3J). However, when we stimulated Target neurons in a part of the VR environment without landmarks (n = 4 mice), we observed an increase in activity that shifted away from the stimulation-paired location (Fig. S3K, L). This suggests that the presence of a landmark may help tether stimulation induced activity to specific positions in the VR.

Outside of the stimulation-paired landmark, Target neuron activity was decreased in post-stimulation trials and recovered on Day 3 (Fig. S3M, N). The decrease in activity outside the stimulation-paired landmark was not correlated with the increase in activity around the stimulation-paired landmark (rho = 0.06, p = 0.34, n = 250 Target neurons). We also did not discover other factors that allowed us to predict which Target neurons would be successfully biased (Fig. S3O-R). As in RSC, these results show that stimulation-induced effects are variable in V1 L2/3 neurons. In addition, it is likely that stimulation leads to short term changes in Target neuron excitability or inhibitory input. Together, these data highlight a plasticity mechanism that generalizes across L2/3 of both V1 and RSC, in which targeted stimulation during initial VR exploration can rapidly bias neural activity to specific visual cues.

We then asked if, similar to RSC, rapid plasticity in V1 was restricted to when mice explored VR for the first time. In contrast to V1 Target neurons in naïve mice, we did not observe a significant increase in activity around the stimulation-paired landmark in post-stimulation trials in familiar mice (Fig. 3G-I, S4A-C, 115 Target neurons, median = 28, range = 12-17 Target neurons per landmark, 27-31 Target neurons per mouse, with similar stimulation efficiency in familiar and naïve Target neurons, Fig. S4D, E). On Day 3, we observed a small increase in activity in Target neurons (Fig. 3H, J, Fig. S4F, G). As in RSC, the activity outside the stimulation-paired landmark was decreased on post-stimulation trials on Day 2, but recovered on Day 3 (Fig. S4H, I). This short-term decrease in activity was consistent across regions and experience, raising the possibility that stimulation can transiently decrease activity levels in stimulated neurons, possibly via a decrease in excitability or an increase in inhibitory input.

In parallel to the similar effect of experience on stimulation-induced plasticity between RSC and V1, spatial representations in the left-out population in V1 were initially less stable in naïve mice (Fiser et al. 2016; Pakan et al. 2018) and rapidly stabilized by Day 3 of imaging, while spatial representations remained stable throughout the experiment in familiar mice (Fig. 3K-M). These findings suggest that in familiar environments, the process allowing for strong and rapid plasticity in L2/3 of V1 or RSC is diminished. However, this does not preclude the presence of a slower acting plasticity mechanism induced by stimulation, given the small increase in landmark-surrounding activity in Target neurons in both RSC and V1 on Day 3.

### Novel VR context is sufficient to partly reinstate rapid plasticity

When mice first explore visual VR, the novelty of the experience encompasses more than the visual cues of the VR environment. For example, mice also first experience that running on the treadmill causes visual flow. To isolate the effect of environmental novelty related to visual cues, we exposed mice that had undergone the familiar environment condition (including stimulation trials) to a novel VR environment. Mice rapidly adapted to the novel environment (Fig. S5A) and in the left-out populations of both RSC and V1, representations for the novel environment stabilized quickly (Fig. S5B, C).

In this novel VR environment, we then performed the same three day experiment in L2/3 of RSC or V1 (Fig. 4A). We stimulated Target neurons in L2/3 of RSC (303 stimulated and tracked neurons, 7 mice, median = 22, range = 13-28 Target neurons per landmark, 30-53 Target neurons per mouse) or V1 (192 stimulated and tracked neurons, 5 mice, median = 19, range = 14-25 Target neurons per landmark, 34-40 Target neurons per mouse). In RSC, we did not observe a significant increase in activity around the stimulation-paired landmark in the Target neurons in trials following the stimulation trials (Fig.4 B-D, Fig. S5D, E). However, we did observe a significant increase on Day 3 (Fig. 4E, Fig. S5F-I). In V1, the change in the activity of Target neurons around the stimulation-paired landmark was significant but small in trials following the stimulation trials (Fig. 4F-J, Fig. S5J-L). On Day 3, activity was significantly increased around the stimulation-paired landmark (Fig. 4K, S5L-O). This was due to both Target neurons that had increased activity on Day 2 post-stimulation trials (Example in Fig. 4F i) and Target neurons that had increased activity only on Day 3 (Example in Fig. 4F ii, iii, Fig. S5O). These results suggest that stimulation triggers both rapid (Day 2) and slower-acting (Day 3) plasticity changes when novelty is limited to the VR environment alone.

**Figure 4:**
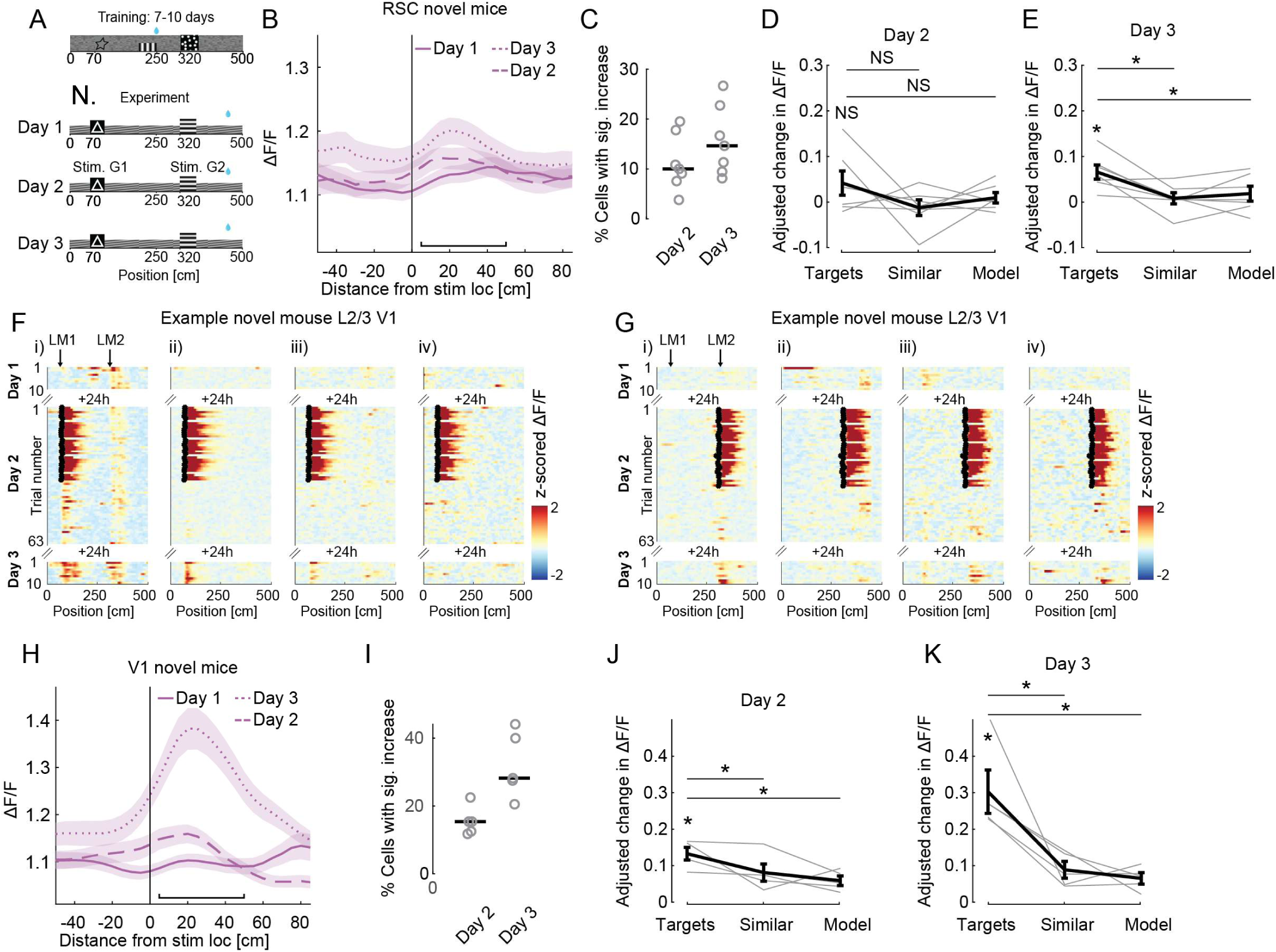
A visually novel environment increases stimulation-evoked plasticity. A. Schematic for experiments in a novel VR environment (‘novel’ mice). Mice are familiarized with one visual VR environment, and the experiment is conducted in a second VR environment, with different landmarks and visual textures, and a different reward location. B. Mean spatially binned activity around the stimulation-paired landmark of all Target neurons in L2/3 RSC of novel mice. Data from LM1 and LM2 Target neurons are combined (n = 303 Target neurons, 7 mice, range = 30-53 Target neurons per mouse). Day 1 (solid line), Day 2 after stimulation trials ended (long dash) and Day 3 (short dash). Shading indicates SEM. Black horizontal line: landmark-surrounding region used for quantification. C. The fraction of Target neurons per mouse (n = 7 mice) with a significant increase in landmark-surrounding activity on Day 2 and Day 3, relative to activity on Day 1 (as in Fig. 1I). Grey circles: average fraction per mouse. D. No significant change in activity around the stimulation-paired landmark between Day 1 and Day 2 for Target neurons (left), compared to Similar and Model neurons (as in Fig. 1J) (Targets vs. Similar: p > 0.05, Targets vs. Model: p > 0.05, Targets vs. 0: p > 0.05, one-sided Wilcoxon signed-rank test, 7 mice). Black line indicates mean ± SEM, gray lines indicate mean change per mouse. E. Same as in (D), but for the difference in activity between Day 1 and Day 3. There was a significant increase in activity around the stimulation-paired landmark in Target neurons. (Targets vs. Similar: p ≤ 0.02, Targets vs. Model: p ≤ 0.04, Targets vs. 0: p ≤ 0.01, one-sided Wilcoxon signed-rank test, n = 7 mice). F. Spatially binned activity of four co-recorded Target neurons from V1 of an example novel mouse. Target neurons are from group 1, with stimulation paired only to landmark 1 (LM1) on Day 2. Activity around LM1 is significantly increased in Target neurons on the left (i), but unchanged in iv (i: p < 0.025, iv: p > 0.05 one-tailed Wilcoxon signed-rank test, comparing LM1 surrounding activity on 10 post-stimulation trials to activity on Day 1). Examples in ii) and iii) are unchanged on Day 2, but show a significant increase on Day 3 (ii, iii: p < 0.025 one-tailed Wilcoxon signed-rank test, comparing LM1 surrounding activity on Day 3 trials to activity on Day 1). Arrows on top indicate the position of stimulation-paired landmarks. Small black x’s mark the onset of stimulation. G. As in (F), but for group 2 Target neurons from a different example mouse. H. Same as in (B), but for Target neurons in V1 of novel mice (n = 192 Target neurons, 5 mice, range = 34-40 Target neurons per mouse). I. Same as in (C), but for Target neurons in V1 of novel mice (n = 5 mice). J. Change in adjusted landmark-surrounding activity per mouse between Day 1 and Day 2 for Target neurons (left), compared to Similar and Model neurons. Change in activity in Target neurons is significantly larger than in Similar and Model neurons. (Targets vs. Similar: p ≤ 0.03, Targets vs. Model: p ≤ 0.03, Targets vs. 0: p ≤ 0.03, one-sided Wilcoxon signed-rank test, n = 5 mice). Black line indicates mean ± SEM, gray lines indicate mean change per mouse K. Same as in (J), but comparing activity between Day 1 and Day 3 (Targets vs. Similar: p ≤ 0.03, Targets vs. Model: p ≤ 0.03, Targets vs. 0: p ≤ 0.03, one-sided Wilcoxon signed-rank test, n = 5 mice)

### Layer 5 neurons in RSC maintain plasticity independent of environmental familiarity

Our data show that stimulation-induced plasticity in L2/3 of both V1 and RSC is limited to a subset of neurons and strongly modulated by experience. We next investigated if similar effects could be observed in layer 5 (L5) RSC neurons, which differ from L2/3 neurons in their intrinsic and functional properties (Kurotani et al. 2013; Fischer et al. 2020; Mao et al. 2017; Sempere-Ferràndez, Andrés-Bayón, and Geijo-Barrientos 2017; Sullivan et al. 2023), as well as how neuromodulation influences their plasticity dynamics (Chubykin et al. 2013; He et al. 2015). To gain optical read-write access to L5 of RSC, we crossed the *Ai228* mice to the L5 Cre driver *Rbp4-Cre* (Drinnenberg et al. 2024). *Ai228;Rbp4-Cre* mice expressed GCaMP6m and ChRmine-oScarlet in L5 neurons (Fig. 5A, B). We then used the same approach in L5 of naïve mice (n = 3 naïve mice) and familiar mice (n = 5 familiar mice). Neurons were stimulated on Day 2 for 30 trials (0.75 second stimulation at 47 Hz) with three no-stimulation trials interleaved (trials 8, 18 and 28). Stimulation trials were followed by at least 10 non-stimulation trials and activity was recorded from the same neurons on Day 3 to assess longer lasting changes. This allowed us to investigate if activity can be biased towards stimulation-paired locations in a manner that is similar to our observations in superficial layers of RSC and if familiarity with the environment plays a role in modulating plasticity.

**Figure 5.**
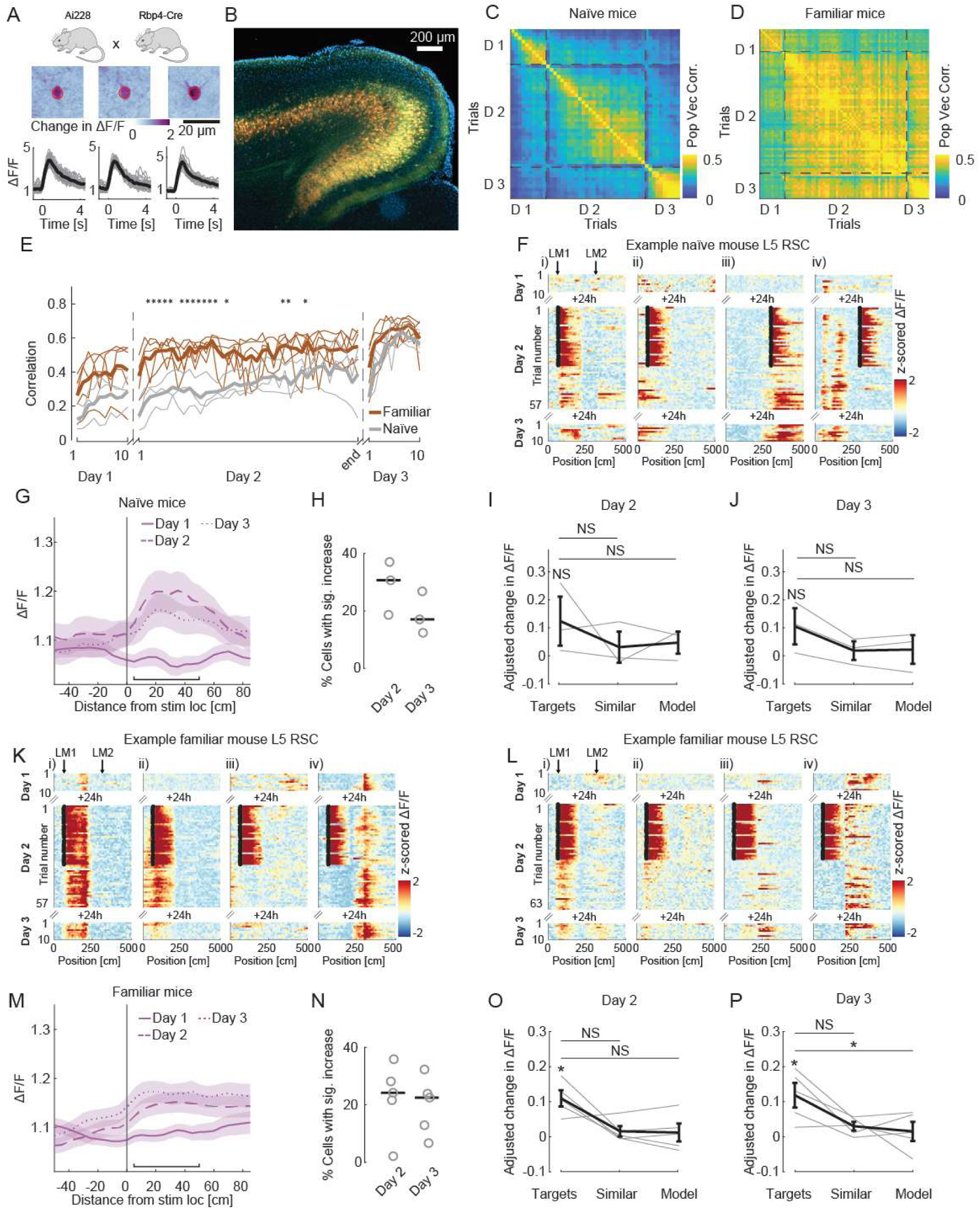
Stimulation evoked plasticity in layer 5 does not depend on experience. A. Top: We used *Ai228;Rbp4-Cre* mice to restrict the expression of GCaMP6m and ChRmine-oScarlet to L5 neurons. Middle: ΔF/F images of the stimulation-evoked activity, centered around 3 stimulation targets in L5 of RSC. Bottom: Single trial (gray lines) and trial-averaged (black line) stimulation-evoked activity of the same neurons (n = 30 stimulation trials). B. Fluorescence image of an example coronal section of an *RBP4-cre;Ai228* mouse showing GCaMP6m (green) and oScarlet (red) co-expression in Layer 5 of cortex. Scalebar: 200 µm C. Spatial similarity in non-stimulated L5 neurons across trials in naïve mice. Heatmap shows population vector correlation of position-stable neurons, comparing activity between all trials within and across three experimental days (D1, D2, D3). Dashed lines indicate day boundaries. (n = 3 mice, median = 157, range = 138-194 position-stable left-out neurons). D. Same as (C), but spatial similarity matrix for all familiar mice (n = 5 mice, median = 195, range = 146-240 spatially stable left-out neurons). E. Correlation of spatial activity patterns of left-out neurons between Days 1 and 2 and Day 3 in familiar (brown) and naïve (gray) mice. Higher correlation indicates greater similarity to Day 3. Thick lines: mean; thin lines: individual mice. *p < 0.05, **p < 0.01 (Mann-Whitney U test, n = 8 naïve mice, n = 4 familiar mice). Day 2 data is interpolated to accommodate varying trial numbers (Methods). F. Spatially binned activity of four co-recorded L5 Target neurons from an example naïve mouse. Target neurons i and ii are from group 1, with stimulation paired to LM1, Target neurons iii and iv are from group 2, with stimulation paired to LM2 on Day 2. Activity around stimulation-paired landmark is significantly increased in i and iii (i, iii: p < 0.025, ii, iv: p > 0.05 one-tailed Wilcoxon signed-rank test, comparing landmark-surrounding activity on 10 post-stimulation trials to activity on Day 1). Arrows on top indicate the position of stimulation-paired landmarks. Small black x’s mark the onset of stimulation. G. Mean spatially binned activity around the stimulation-paired landmark of all L5 Target neurons in RSC of naïve mice (n = 94 Target neurons, 3 mice, range = 26 - 35 Target neurons per mouse), for the three imaging sessions (Day 1: solid, Day 2: long dash, Day 3: short dash). Shading indicates SEM. Black horizontal line: ‘landmark-surrounding’ region used for quantification. H. The fraction of Target neurons per mouse (n = 3 mice) with a significant increase in landmark-surrounding activity on Day 2 and Day 3, relative to activity on Day 1. I. Change in landmark-surrounding activity per mouse between Day 1 and Day 2 for Target neurons is not statistically different from activity in Similar and Model neurons (Targets vs. Similar: p > 0.05, Targets vs. Model: p > 0.05, Targets vs. 0: p > 0.05, one-sided Wilcoxon signed-rank test, n = 3 mice). Black line indicates mean ± SEM, gray lines indicate mean change per mouse. Note that the relatively small number of animals makes the statistical test difficult to interpret. See also Fig. S6B. J. Same as in (I), but comparing activity between Day 1 and Day 3. See also Fig. S6E. K. Spatially binned activity of four co-recorded L5 Target neurons from an example familiar mouse. Target neurons are from group 1, with stimulation paired to only landmark 1 (LM1) on Day 2. Activity around LM1 is significantly increased in Target neurons on the left (i and ii), but unchanged in iii and iv (i, ii: p < 0.025, iii, iv: p > 0.05 one-tailed Wilcoxon signed-rank test, comparing LM1 surrounding activity on 10 post-stimulation trials to activity on Day 1). Arrows on top indicate the position of stimulation-paired landmarks. Small black x’s mark the onset of stimulation. L. Same as in (K), but for four Target neurons from a different mouse. Activity around LM1 is significantly increased in Target neurons on the left (i and ii), but unchanged in iii and iv (i, ii: p < 0.025, iii, iv: p > 0.05 one-tailed Wilcoxon signed-rank test, comparing LM1 surrounding activity on 10 post-stimulation trials to activity on Day 1) M. Mean spatially binned activity around the stimulation-paired landmark of all L5 Target neurons in RSC of familiar mice (n = 205 Target neurons, 5 mice, range = 25 - 53 Target neurons per mouse). N. The fraction of Target neurons per mouse (n = 5 mice) with a significant increase in landmark-surrounding activity on Day 2 and Day 3, relative to activity on Day 1. Grey circles: average fraction per mouse. O. Change in landmark-surrounding activity per mouse between Day 1 and Day 2 for Target neurons is not statistically different from activity in Similar and Model neurons, but larger than 0 (Targets vs. Similar: p = 0.06, Targets vs. Model: p = 0.06, Targets vs. 0: p ≤ 0.03, one-sided Wilcoxon signed-rank test, n = 5 mice). Black line indicates mean ± SEM, gray lines indicate mean change per mouse. P. Same as in (O), but comparing activity between Day 1 and Day 3 (Targets vs. Similar: p = 0.06, Targets vs. Model: p ≤ 0.03, Targets vs. 0: p ≤ 0.03, one-sided Wilcoxon signed-rank test, n = 5 mice).

We found that spatial representations in left-out L5 neurons were initially unstable and stabilized with experience in naïve mice (Fig. 5C), and were more stable in familiar mice (Fig. 5D, E). In naïve mice, pairing visual landmarks with targeted stimulation on Day 2 led to a lasting increase in landmark-surrounding activity in a subset of Target neurons in L5 (Fig. 5F-J, S6A-F), similar to our observations in L2/3 of both RSC and V1. In contrast to L2/3, where we observed little stimulation-induced plasticity in familiar mice, targeted stimulation on Day 2 led to a lasting increase in landmark-surrounding activity in L5 Target neurons of familiar mice (Fig. 5K-P, S6G-L). Our data show that in L5 neurons of RSC, activity can be biased towards stimulation-paired landmarks, like in L2/3. However, experience does not appear to influence stimulation-induced plasticity in deeper RSC layers as strongly as it does in superficial layers.

## DISCUSSION

Cortical circuits continuously balance plasticity during new learning with the stability required to maintain established representations. How the dynamics of this balance differs between different brain regions, layers, and over experience remain incompletely understood. Using precise all-optical manipulations in behaving mice, we investigated these questions in RSC and V1 during navigation in VR environments. Our findings reveal a layer-specific mechanism governing such a balance between plasticity and stability. Superficial layers of both RSC and V1 exhibited rapid plasticity, in novel but not familiar environments, while layer 5 retained plasticity regardless of environmental novelty. Modifications in neural activity emerged within a single session, and persisted across days, allowing us to investigate cortical plasticity dynamics during ongoing behavior.

We observed remarkably similar plasticity between RSC and V1, including the proportion of persistently biased Target neurons and the strong modulatory effect of environmental familiarity. In addition, the spatial representations in both RSC and V1 were broadly similar, rapidly stabilizing early in learning and remaining stable in familiar environments. These similarities are consistent with transcriptomic evidence suggesting conserved cell type architectures between agranular RSC and primary visual cortex (Sullivan et al. 2023; Tasic et al. 2018), raising the possibility of conserved plasticity mechanisms across cortical regions. For example, specific subtypes of CamKIIα-positive excitatory neurons might be more susceptible to stimulation induced plasticity in both regions (Tasic et al. 2018; O’Toole, Oyibo, and Keller 2023; Schneider-Mizell et al. 2025). This could help explain why only a fraction of Target neurons significantly increased their activity after stimulation. Future work could examine if specific subtypes of L2/3 neurons preferentially exhibit plasticity, which could provide further insight regarding the potential functional role of different classes of excitatory neurons in L2/3.

Our data suggest that plasticity in L2/3 is modulated by novelty. One mechanism by which novelty has been proposed to regulate plasticity is via neuromodulatory systems. The locus coeruleus (LC) is strongly activated by unexpected or novel stimuli (Takeuchi et al. 2016; Jordan and Keller 2023; Breton-Provencher et al. 2022), and LC activity has been implicated in long-term potentiation (LTP) and memory formation via the release of both dopamine and norepinephrine (Takeuchi et al. 2016). Prior work has shown that norepinephrine enhances LTP (He et al. 2015; Hong et al. 2022) and visuomotor plasticity (Jordan and Keller 2023) in L2/3 of V1. Furthermore, novel and unpredicted stimuli activate vasoactive intestinal peptide (VIP) interneurons (Furutachi et al. 2024; Garrett et al. 2020, 2023; Attinger, Wang, and Keller 2017; Tamboli et al. 2024). VIP interneurons have been shown to be potent regulators of plasticity in the brain (Williams and Holtmaat 2019; Canto-Bustos et al. 2022; Neubrandt, Lenkey, and Vervaeke 2025), potentially via disinhibition of distal dendrites of pyramidal cells. An experience-dependent decrease in LC activity or reduced activity of VIP neurons could thus constrain superficial cortical plasticity in familiar environments.

Another potential neuromodulator that may play a role in experience-dependent cortical plasticity is acetylcholine, which promotes LTP in L5 but not L2/3 of V1 (Chubykin et al. 2013; He et al. 2015). As acetylcholine has been generally associated with arousal and locomotion, a baseline cholinergic signal in familiar environments could be sufficient for plasticity in L5 irrespective of experience. Thus, our findings, together with previous work, suggest that neuromodulation could provide a mechanism for protecting stable L2/3 representations of familiar environments from excessive plasticity, while allowing persistent plasticity in L5 for continuous updates of relevant contextual information.

Both the increase in activity at the stimulation-paired location, and the decrease in activity at non targeted location manifested rapidly, with changes observable after 10-20 pairings. These observations are reminiscent of BTSP in CA1, where stimulation induced changes to place cell activity can occur on a similarly rapid timescale. However, a key feature of BTSP in the hippocampus is the broad temporal window for plasticity. In CA1, synapses active seconds before or after the plasticity-inducing event can be potentiated, resulting in backward or forward shifting of place cell activity along the track, a feature we did not observe in our data. Instead, the increase in activity we observe appears to be more strongly anchored to the stimulation-paired location, suggesting a tighter temporal window for cortical plasticity. Nevertheless, it remains possible that stimulation leads to calcium-events in the dendrites of Target neurons, which then recruit molecular mechanisms similar to those involved in BTSP. Finally, we also observed an overall decrease in stimulated neurons outside the stimulation-paired landmark. This argues against an overall increase in cellular excitability in Target neurons as the primary mechanism for the increase in activity at the stimulation-paired location and instead supports the idea that plasticity reflects synapse specific changes.

In summary, our findings reveal a layer-specific mechanism for controlling cortical plasticity based on environmental familiarity. This experience-dependent modulation of rapid plasticity in superficial layers, contrasted with persistent plasticity in deep layers, offers a potential circuit level framework for how the brain adaptively encodes new environments while preserving stable representations of the familiar world. These results thus provide a key foundation for future work focused on dissecting the precise interplay of neuromodulators and cell-type specific circuits driving these distinct layer-dependent dynamics, as well as determining how stimulation induced plasticity relates to naturally occurring plasticity.

## Acknowledgments

We thank Adriana Diaz for help with animal husbandry and colony maintenance. We thank all members of the Giocomo lab for feedback and input. L.M.G. and K.D. are Howard Hughes Medical Institute Investigators. Funding was provided by the US National Institutes of Health grants BRAIN Initiative U19NS118284 (to K.D. and L.M.G.), P50 DA042012 (to K.D. and L.M.G.), 1R01MH126904-01A1 and R01MH130452 (to L.M.G.), The Vallee Foundation (to L.M.G.), The James S. McDonnell Foundation (to L.M.G.), The Simons Foundation 542987SPI (to L.M.G.), the Gatsby Foundation (to K. D.) and the Swiss National Science Foundation P400PB_191076 (to A.A)

## Author Contributions

A. Attinger, A. Drinnenberg and L.M. Giocomo conceptualized the study. A. Attinger, A. Drinnenberg, La’Akea Siverts, Tanya Daigle, Bosiljka Tasic, Hongkui Zeng, Sean Quirin, Karl Deisseroth and L.M. Giocomo contributed to the methodology. A. Attinger, A. Drinnenberg and C. Dong performed the investigation. A. Attinger performed the visualization. A. Attinger, L.M. Giocomo and K. Deisseroth acquired funding. A. Attinger and L.M. Giocomo conducted project administration. L.M. Giocomo and K. Deisseroth supervised the project. A. Attinger and L.M. Giocomo wrote the original draft of the manuscript. All authors reviewed and edited the manuscript.

## Declarations of Interests

K.D. is a founder and scientific advisor for Maplight Therapeutics and Stellaromics and a scientific advisor to RedTree LLC and Modulight. All materials and protocols are freely available to nonprofit institutions and investigators.The other authors declare no competing interests.

## Data Availability

Upon publication, data will be made available on a publicly accessible data sharing site (e.g. figshare, DANDI, Mendeley).

## Code Availability

Upon publication, custom scripts for analyzing the data will be available on GitHub.

## Methods

### Subjects

All experimental procedures were approved by the Institutional Animal Care and Use Committee at Stanford University School of Medicine. For layer 2/3 experiments, we used a cohort of n = 23 double-positive *Ai228;Camk2a-CreERT2* mice (9 male, 14 female) generated by crossing *Camk2a-cre/ERT2* mice (Jackson Laboratory strain number: 012362) with *Ai228* mice (Drinnenberg et al. 2024). Mice were aged 9.5 to 21.5 weeks (weighing 18−31 g) at the time of surgical implantation. Cre recombination was induced by administering three doses of tamoxifen (100 μL of 20 mg/mL solution in corn oil; Sigma-Aldrich, T5648-1G) via intraperitoneal injection, spread over 5 days, starting 2 weeks prior to surgery. For layer 5 experiments, n = 5 double-positive *Ai228;Rbp4-Cre* mice (3 male, 2 female, aged 8.5 to 12 weeks at surgery) were generated by crossing *Rbp4-Cre* (Tg(Rbp4-cre)KL100Gsat, No. 031125-UCD, MMRRC) (L. Luo)) with *Ai228* mice. All mice had ad libitum access to food and water before surgery and ad libitum access to food throughout the experimental period.

### Surgery

During the imaging window implantation procedure, mice were anesthetized via an intraperitoneal injection of ketamine (85 mg/kg) and xylazine (8.5 mg/kg). Anesthesia was maintained throughout the procedure with inhaled 0.5−1.5 % isoflurane in 0.5 L/min oxygen, delivered via a standard isoflurane vaporizer. To minimize inflammation, mice received two intraperitoneal injections of dexamethasone (2 mg/kg), one 24 hours and one 2 hours prior to surgery. To provide analgesia, Rimadyl (5−10 mg/kg) was injected i.p. immediately before surgery.

A 5 mm craniotomy was performed over the right posterior cortex, extending across the midline, targeting the retrosplenial cortex (RSC) and primary visual cortex (V1). A pneumatic dental drill (NSK Pana-Max PAX-TU M4) equipped with a carbide bur (Meisinger, HM1-005-HP Round Operative Carbide Bur US#1 / 4, 0.5 mm ∅, HP) was used, ensuring the dura remained intact. A 5 mm circular cover glass (Warner Instruments: 64-0700) was then secured flush with the skull using cyanoacrylate adhesive (Loctite Super Glue). Finally, a custom-fabricated titanium head-plate was affixed to the skull using Metabond dental acrylic, dyed black with India ink or black acrylic powder for light shielding.

Post-surgically, mice received 1 mL of saline and 10 mg/kg of Baytril via subcutaneous injection and were placed on a warming blanket for recovery. Mice were monitored for several days, with additional Rimadyl and Baytril administered as needed to address signs of discomfort or infection. A recovery period of 1−2 weeks was allowed before initiating water restriction and behavioral training.

### All-Optical Two-Photon Holographic Experiments

#### Microscope System and Imaging

Our custom-built all-optical two-photon (2P) microscope system integrated a holographic stimulation pathway with a commercial 2P microscope (Bruker Ultima 2P plus, controlled by PrairieView software). This system was based on the multiSLM setup previously described by (Marshel et al. 2019), with key modifications including an increased imaging field-of-view and the incorporation of an electro-tunable lens for rapid axial scanning and multi-plane imaging.

Calcium transients from GCaMP6m were imaged at 920 nm using a tunable femtosecond laser (Coherent Discovery with TPC). Imaging laser power was dynamically increased by approximately 3-4% for the intermediate imaging plane, and 6-8% for the deepest imaging plane via the settings in the PrairieView software. Typical power ranges measured below the objective during scanning were: 15 − 25 mW in L2/3 RSC and V1 and 25 - 30 mW in CA1 for *Ai228;CamK2a-CreERT2* mice; and 30−40 mW in L5 for *Ai228;Rbp4-Cre* mice. Imaging was performed using a 16×/0.8 NA objective (Nikon) immersed in diluted ultrasound gel.

For L2/3 experiments, data were acquired at a resolution of 1024×1024 pixels within a 1×1 mm^2^ field-of-view (FOV) at 5 Hz per plane. For L5 experiments, the FOV was approximately 0.75×0.75 mm^2^, with data acquired at 512×512 pixels resolution at 10 Hz per plane. An electro-tunable lens enabled fast, sequential acquisition of three planes, separated by 40 μm (30 μm for CA1 experiments).

#### Holographic stimulation

Stimulation light was generated by a fixed-wavelength (1035 nm) femtosecond laser (Coherent Monaco 1035-80-60) operating at a 2 MHz pulse repetition rate. Holograms were computed from 3D target coordinates using an algorithm implemented in MATLAB, as described in Marshel et al., 2019 (Marshel et al. 2019), and projected via a spatial light modulator (Boulder Nonlinear Systems / Meadowlark Optics macroSLM, 1536×1536 pixels with temperature control).

Each hologram was scanned in a 7 μm diameter spiral (5 rotations in 2 ms) across targeted neuron somata using a pair of galvo mirrors. This spiral scan was repeated 31 times at 32 Hz for L2/3 experiments and 31 times at 46 Hz for L5 experiments. To compensate for light scattering at different depths, an amplitude term weighted by the function √(e^(z/ρ)) with scattering length ρ = 250 µm was added to each target in the hologram, where z was the offset of lower imaging planes relative to the uppermost plane. Neurons in L2/3 were stimulated using peak power in the range of 4.0 mW to 5.0 mW per cell (time averaged power: 0.26 mW - 0.33 mW), 6.5 mW to 7 mW per cell in L5 (time averaged power: 0.61 mW to 0.66 mW), and 5.0 mW to 6.0 mW per cell in CA1 (time averaged power: 0.33 mW to 0.39 mW). Stimulation parameters were empirically determined through pilot experiments (data not shown) to achieve robust neuronal activation within a 1-second window. Stimulation caused a 2 ms contamination of the GCaMP signal, affecting approximately 10 lines (20 lines for 10 Hz imaging) per single stimulation event. To mitigate this artifact, the stimulation frequency was optimized to prevent overlapping artifact regions in consecutive images within the same plane. Affected scan lines in images were subsequently replaced by the average of the leading and trailing frames during post-processing.

Optical aberrations of the optogenetic beam were corrected by scanning the optogenetic galvo pair and forming an image using a bulk fluorescent slide. Zernike polynomials (OSA/ANSI single-indexed modes 3−20) were sequentially applied with varying weights to the SLM phase mask, generating performance curves used to select values that maximized image intensity (i.e., minimized optical aberration), as previously described (Marshel et al. 2019). For stimulation during VR behavior, individual stimulation bouts were triggered by the VR software (see Methods section below on Virtual Reality), and the gating signal controlling individual pulses was recorded concurrently with imaging data for precise synchronization.

#### Alignment of Stimulation and Imaging Paths

Prior to each stimulation experiment, precise alignment was ensured by burning a predefined 3D pattern of holes into a fluorescent slide using the stimulation laser. The centers of these holes were then manually identified in the 2P image across multiple planes. By comparing the coordinates of the holes to the intended location of the calibration pattern, we identified and corrected for lateral (1−3 pixel) and axial (1−4 μm) shifts between the imaging and stimulation paths.

#### Online Motion Correction

At the start of each experimental series, a reference image stack (21 planes, Δz=1 μm, spanning ± 10 μm around the center imaging plane, averaged over 50 frames) was acquired. A custom MATLAB script, interfacing with PrairieView, performed online motion correction by registering live images to the center plane of the reference stack using a cross-correlation-based algorithm in real time. This approach was also used for day-to-day alignment of the field of view, after approximate realignment by prominent landmarks such as blood vessels.

#### Virtual Reality (VR) Setup, Design and Behavior

All VR tasks were developed using custom Python code based on the open-source Panda3D game engine (https://www.panda3d.org/). The VR environment was projected using a DLP LightCrafter projector (TI CLPDLCR2010EVM), adapted from https://github.com/HarveyLab/mouseVR, and refreshed at a frame rate of 50 Hz. The green LED of the projector was disabled to minimize crosstalk with the imaging system.

The VR behavioral system incorporated a 6-inch diameter fixed-axis cylindrical treadmill. Treadmill rotation was recorded via a rotary encoder (Yumo, 1024 P/R), with data processed and transmitted to the VR software via an Arduino UNO microcontroller running custom firmware. A capacitive lick port, comprising a feeding tube (Kent Scientific) and an 18G gavage needle (2 mm tip, https://www.petsurgical.com/animal-feeding-gavage-needle-reusable-straight-curved/) wired to a capacitive sensor (Adafruit: AT42QT1010), detected licks and delivered 5 % (weight/volume) sucrose water reward via a gravity solenoid valve (Cole Palmer), calibrated to deliver 6−10 μL per reward.

#### Synchronizing VR and Imaging

Synchronization between VR and imaging data was achieved using a digital signal generated by the VR software via an Arduino UNO. This signal alternated between ’low’ and ’high’ on consecutive VR frames and was recorded as an additional input on the imaging computer, enabling precise alignment of VR data timestamps with imaging frame timestamps.

#### Behavioral Tracking

VR software recorded mouse position, running speed, and lick sensor voltage at the VR frame rate (50 Hz). All VR data were resampled to the imaging frame times using linear interpolation during data analysis. Running speed was smoothed using a normalized Gaussian kernel with a width of 5 frames and a standard deviation of 1.

#### Behavioral Training

Following a 1-week post-surgical recovery period, mice underwent 2 weeks of water restriction and behavioral training, with 1-hour daily sessions on the setup. Initially, mice were trained to run on the cylindrical treadmill in darkness, receiving rewards at very short inter-reward distances. This distance was progressively increased to 8 m. Reward size was approximately 5 μL, and inter-reward distance was adjusted automatically to maintain approximately 200 rewards per hour, up to a maximum of 8 m.

#### VR Environment Design

The VR environment generally consisted of a black floor and patterned walls. The gain on the wheel was adjusted such that 1 position unit corresponded to 1 cm. Environments were 500 cm long, except for experiments in 4 V1 mice, where the environment was extended to 550 cm, incorporating an additional segment with only visual flow cues. Upon reaching the end of the environment, the screen was grayed out for 3−8 seconds (drawn from a uniform distribution for each trial) before the position of the mouse was reset to the starting position for a new trial. For “naïve” and “familiar” environments, walls were patterned with rectangles of randomized grayscale values to provide optic flow cues. These environments also contained static cues at positions 50, 250, and 400 cm, each consisting of a gray narrow cylinder rotated uniquely to provide distinct spatial cues. An additional cue, a sine grating replacing the left wall cue (contralateral to the recording side) for 60 cm, was present in both naïve and familiar environments in the region immediately preceding the reward location. Reward was delivered unconditionally at position 250 cm (450 cm in the novel environment). If mice licked within a 10 cm zone preceding the reward location, reward was delivered early.

##### Stimulation-paired landmarks

Naïve and familiar environments included two transient landmarks that appeared for 1 second when mice reached specific positions (70 cm for landmark 1, 320 cm for landmark 2). These landmarks moved with the environment based on mouse speed and were not necessarily visible for the entire 1 second duration. They comprised flat elements textured with unique patterns (e.g., a single black star on a white background for landmark 1, black dots on a white background for landmark 2) and replaced the left wall pattern. For the novel environment, stimulation-paired landmarks appeared at 70 cm (landmark 1) and 320 cm (landmark 2) and consisted of a white triangle on a black background (landmark 1) or a black outline of a star on a gray background (landmark 2).

### Imaging and Stimulation Experiments Across Days

#### Naïve Experiments

Following behavioral training, mice underwent three experimental days (Fig. 1A). The day prior to the experiment Day 1, mice were screened with the 2P microscope to identify a suitable imaging location. On Day 1, a reference stack was recorded for online and day-to-day alignment. We then recorded neuronal activity for 8−10 minutes as mice ran in darkness, receiving rewards every 800 cm (average 10.7 ± 1.5 rewards, mean ± SEM). This dark session ensured sufficient data for registration and automatic ROI extraction (see below) and allowed us to further characterize neural activity. A VR session immediately followed, consisting of 10 traversals of the environment described above, with automatic reward delivery. After the VR session, mice continued running in darkness for an additional 10 minutes, during which oScarlet expression was briefly imaged at 980 nm (500 frames per plane). On Day 2, after returning to the same FOV and ensuring optimal alignment, mice traversed the same environment an average of 53 times (median = 53, min = 45, max = 68). Stimulation of target neurons was position-triggered: when mice passed 70 cm (landmark 1) or 320 cm (landmark 2), a signal was sent via an Arduino UNO to the stimulation system. In 4 out of 7 V1 ’naïve’ mice, an additional stimulation trigger was sent at 480 cm, activating a third ensemble. Stimulation commenced at the beginning of Day 2 and continued until each group of target neurons was stimulated 30 times. In all but 2 naïve mice, stimulation trials were interleaved with three non-stimulation trials (trials 7,14, 21). After the final stimulation trial, mice ran an additional average of 19 trials (median = 19, min = 10, max = 38) during which landmarks were displayed but no stimulation was triggered. Unless otherwise noted, the first 10 post-stimulation trials were used for Day 2 analysis. Following VR experiments, an additional 5−10 minutes of dark running was recorded. On Day 3, mice traversed the environment for 10 or more trials while we recorded activity in the same population of neurons. Only the first 10 trials were used for analysis.

#### Familiar Experiments

The structure of familiar experiments was identical to naïve experiments, except that mice first underwent the ’naïve’ protocol. Subsequently, they were trained for an additional 7-10 sessions in the VR environment, 1 hour per session. On Day 2, mice performed an average of 56 trials (median = 56, min = 45, max = 67), with an average of 24 post-stimulation trials (median = 24, min = 10, max = 33). In cases where familiar mice underwent stimulation experiments in the ‘naïve’ protocol, we moved the FOV within the brain region, or alternated brain regions (V1 and RSC) to ensure that different populations of neurons were stimulated.

#### Novel Experiments

The structure of the novel experiment was similar, except that on Day 1, we recorded data during a brief dark session, followed by on average 11 trials in the familiar environment (median = 11, min = 5, max = 25). This was followed by 10 trials in the novel environment. On Day 2, mice performed on average 58 trials (median = 58, min = 49, max = 75), with an average 24 post-stimulation trials (median = 24, min = 12, max = 35).

#### Selection of Stimulation Targets

Day 1 imaging data was processed the same day. Target neurons were selected manually using multiple criteria. Putative target neurons for stimulation at landmark 1 (or landmark 2) were identified as either position-stable neurons lacking an activity field at the respective landmark, or as non-position-stable neurons (trial-to-trial spatial correlation < 0.1). Putative target neurons within 20 μm of the imaging field-of-view border were excluded. Using a custom MATLAB tool, putative target neurons in each plane were highlighted on the average 2P images of both the green (GCaMP6m expression) and red (oScarlet expression) channels. Final target neurons were selected from this pool by visually confirming robust co-expression of GCaMP6m and oScarlet. Target coordinates from each plane were then transformed into the SLM coordinate frame, accounting for the z-coordinates of each plane.

#### Quantification and Statistical Analysis

All analyses were performed in MATLAB or Python using custom code. Nonparametric tests or permutation tests were predominantly employed to avoid assumptions about data distributions.

#### Imaging Data Processing

The Suite2P software package (v0.10.3) (Pachitariu et al. 2016) was used for non-rigid xy motion correction and identification of putative cellular regions of interest (ROIs). Manual curation was performed to eliminate ROIs containing multiple somata or dendrites. Subsequent analysis was performed in MATLAB.

Neuropil contamination was reduced by estimating and subtracting neuropil signals for each ROI, following the method described by Dalgleish et al. (Dalgleish et al. 2020). Neuropil subtraction coefficients were determined using robust regression (MATLAB’s robustfit function with ’bisquare’ weighting) between the 10× downsampled ROI signal and the 10× downsampled neuropil signal. Coefficients were constrained between 0.5 and 1; if regression failed to converge, the median of well-fit coefficients was used. The resulting neuropil-corrected traces were then re-baselined to match the baseline of the original, uncorrected traces. To correct for slow baseline drift, a running baseline (20 s sliding window) was subtracted from the neuropil-corrected calcium traces. The resulting traces were then normalized to their median value to obtain ΔF/F values.

Imaging data was processed for each day separately. Custom code was used (Sosa, Plitt, and Giocomo 2025) to identify cells successfully tracked across imaging sessions based on the intersection over union (IOU) of their ROI pixels. The threshold for ROI matching was algorithmically chosen for each dataset such that the IOU for the best match for an ROI pair was always greater than the IOU for any second-best match. Unless otherwise stated, only neuron ROIs identified on Day 1 and successfully tracked to Day 2 and Day 3 were included in the analysis.

#### Assigning Stimulation Targets to Neurons

To assign stimulation targets to Suite2P-identified neurons, we determined the Euclidean distance from each target location to the closest extracted neuron ROI. If this distance was less than 8 pixels, the neuron ROI was identified as a target neuron. Left-out neurons were defined as neurons with a planar distance greater than 25 μm from all target locations.

For each stimulation bout, the activity time series in the window spanning 0 s to 3 s post-onset of stimulation was extracted for each stimulation target. Stimulation reliability was defined as the mean trial-by-trial correlation of the stimulation-evoked time series. To exclude unreliably targeted neurons (due to poor target placement or insufficient ChRmine expression), target neurons with stimulation reliability <0.25 were excluded from further analysis.

#### Spatially Binned Activity

Spatially binned activity was computed by dividing the position into 5 cm bins for each trial and averaging the ΔF/F for all frames within a bin. Timepoints where the mouse was stationary (running speed <0.05 cm/s) were excluded. Position bins with missing values in individual trials due to high running speed were filled using linear interpolation. Finally, the spatially binned activity was smoothed using a normalized Gaussian kernel with a width of 3 bins and a standard deviation of 0.6.

#### Landmark Surrounding Activity

Landmark-surrounding activity was defined as the average of the spatially binned activity within a 45 cm window, starting 5 cm after the position that triggered the display of the landmark (5 cm to 50 cm). Unless otherwise stated, landmark-surrounding activity of group 1 Target neurons was computed for landmark 1 (where holographic stimulation of group 1 Target neurons occurred), and landmark-surrounding activity of group 2 Target neurons was computed for the region immediately following landmark 2.

#### Fraction of Targets with increased landmark surrounding activity

To determine if a Target neuron exhibited a significantly increased landmark-surrounding activity after stimulation, we compared the 10 post-stimulation trials on Day 2 (or Day 3 trials) to the 10 baseline trials on Day 1 using a one-sided Wilcoxon signed-rank test. A significant increase in activity was defined as p < 0.025. The fraction of significantly increased Target neurons per mouse was calculated as the sum of all group 1 Target neurons with a significant increase and all group 2 Target neurons with a significant increase, normalized by the total number of Target neurons per mouse.

#### Change in Landmark-Surrounding Activity and Adjusted Change in ΔF/F

We first computed the position series or time series around the stimulation-paired landmark 1 for group 1 Target neurons, and landmark 2 for group 2 Target neurons, and averaged across all Target neurons from all mice within a given day. To visualize the change in ΔF/F as a function of distance (or time) from the target location, we calculated the difference between the average time or position series from Day 2 (or Day 3) and Day 1.

To quantify the change in landmark-surrounding activity of Target neurons, we computed the trial-averaged landmark-surrounding activity for each recording day (Day 1: trials 1−10; Day 2: first 10 post-stimulation trials; Day 3: trials 1−10) for Target neurons, Similar neurons, Model neurons, and left-out neurons. The change in activity was defined as the difference between Day 2 (or Day 3) and Day 1 activity. This difference was computed for all Target neurons, Similar neurons, and Model Neurons (see below), and then averaged within each mouse.

To account for overall changes in activity levels across days, we computed the adjusted change in ΔF/F. In each mouse, we calculated the average difference of a size-matched random sample of the activity difference from the left-out neurons (matching sample size to the number of Target neurons in that mouse). By repeating this 1000 times, we generated a distribution of activity differences within the left-out neurons. For each mouse, the mean of this distribution reflected the average change in landmark-surrounding activity in the left-out neurons. To calculate the adjusted change in activity, we subtracted this average left-out difference from the observed difference in Target, Similar, and Model neurons. Note that this adjustment shifts the baseline but preserves the relative differences between Target, Similar, and Model neurons.

We also compared the average change in landmark-surrounding activity between Day 2 (or Day 3) and Day 1 of Target neurons, as well as Similar neurons, to a shuffle distribution. The shuffle distribution was generated as described above, but using as a pool all left-out neurons from all mice of a given experiment and sampling the total number of Target neurons in each iteration.

#### Stimulation-Evoked Change in Left-out Neurons

For each stimulation-paired landmark in each mouse, we calculated the stimulation-induced difference in ΔF/F as a function of time around landmark onset by subtracting the average activity across the first five post-stimulation trials from that of the last five stimulation trials for each left-out neuron. Neurons were then grouped by distance to their respective closest stimulation target, using 20 μm bins, starting from 25 μm. Finally, these distance-binned responses were averaged across all stimulation-paired landmarks and all mice to generate an estimate of a time-dependent and distance-dependent stimulation-induced change in activity in the population of left-out neurons.

#### Spatial Similarity and Spatial Similarity Matrix

Correlation matrices were computed using the spatially binned activity on each trial. For the spatial similarity of single neurons, we computed the pairwise correlation of activity and averaged across all pairs of trials. Neurons were defined as position-stable or place-correlated if the spatial similarity on any of the 3 days was ≥ 0.2. To compute the trial-by-trial spatial similarity of the population, activity of all neurons within the population was concatenated on each trial and used to compute the trial-by-trial correlation for trials within a day and across days. The spatial similarity matrix was then averaged across mice. To accommodate variations in the total number of trials executed on Day 2, the number of trials considered on Day 2 was capped at the minimum number of trials within that dataset. To compare the activity patterns of individual trials on Day 1 and Day 2 to the average activity on Day 3, we concatenated the activity of all position-stable neurons for every trial and computed the correlation between this concatenated activity and the concatenated, trial-averaged activity on Day 3. To compute the spatial similarity of individual trials i on Day 3, we applied a leave-one-out scheme and computed the trial-averaged activity based on all Day 3 trials, excluding trial i.

#### GLM Model Neurons

To compare the activity of Target neurons to model-based predictions, we trained a separate Generalized Linear Model (GLM) for Target neuron activity based on the activity of the left-out neurons. For each target neuron, a regularized Poisson GLM was trained using the activity (ΔF/F) of all co-recorded left-out neurons (MATLAB *lassoglm*, using ’Poisson’ link function, and λ=0.001). These Model neurons served as a comparison to determine whether Target neuron changes exceeded what would be expected from the overall population dynamics alone. To predict Day 2 and Day 3 data, the model was trained on all Day 1 data. To predict Day 1 data, a prediction for the activity during each trial was generated by training the model on the activity during all other trials (10-fold cross-validation, with data from each trial as a separate group). Predicted ΔF/F values > 20 were discarded and replaced using spline interpolation.

#### Similar Neurons

To account for normally occurring changes in neural representations (such as remapping or activity drift), we identified ’Similar’ neurons as an additional comparison group. To identify the most similar neuron from the pool of co-recorded neurons, we computed a distance metric between the baseline (Day 1) activity of Target neurons and left-out neurons for each Target neuron. The distance metric was computed by comparing the trial-averaged spatially binned activity of Target neurons and left-out neurons using the Euclidean distance. The ‘Similar’ neuron was defined as the left-out neuron with the smallest Euclidean distance to the Target neuron. This comparison allowed us to distinguish stimulation-induced changes from spontaneous changes that might occur in neurons with similar baseline properties

#### Linear models

To identify factors that might predict which Target neurons would show stimulation-induced increase in activity, we performed multiple linear regression analysis using baseline (Day 1) neural properties as predictors of the change in landmark-surrounding activity between Day 1 and Day 2.

Predictor variables included: (1) spatial similarity during Day 1 trials, computed as described above, and (2) correlation with running velocity, computed as the Pearson correlation coefficient between each Target neuron’s activity and smoothed running velocity during the initial dark session on Day 1. Running velocity was smoothed using a Gaussian kernel (5 frames width, σ=1). Multiple linear regression was performed using MATLAB’s *fitlm* with default parameters, including an intercept term.

We also tested whether spatial proximity to other stimulated neurons influenced the response magnitude by predicting the change in landmark-surrounding activity using: (1) planar distance to the closest co-stimulated target, and (2) distance to the center of mass of all co-stimulated targets (computed by averaging x and y coordinates of all targets in the same stimulation group). Multiple linear regression was performed using MATLAB’s *fitlm* with default parameters, including an intercept term.

#### Field width and field peak

For each neuron, we identified isolated activity peaks within a defined spatial window. A peak was defined as the maximum activity occurring within a window ranging from -50 to +90 cm around the stimulation-paired landmark. Peaks were excluded if the maximum occurred at the window’s edge or before the onset of the stimulation-paired landmark. The activity value at the peak location (position-binned) was recorded as the peak amplitude of the place field.

Field width was quantified as the number of contiguous spatial bins surrounding the peak that exhibited activity exceeding the mean activity level within the defined spatial window.

We subsequently compared both field width and peak amplitude between two neuron populations: (1) Target neurons that exhibited both an identifiable peak and a significant activity increase from Day 1 to Day 2, and (2) control neurons (left-out population) that showed comparable activity increases.

---

**Figure S1:**
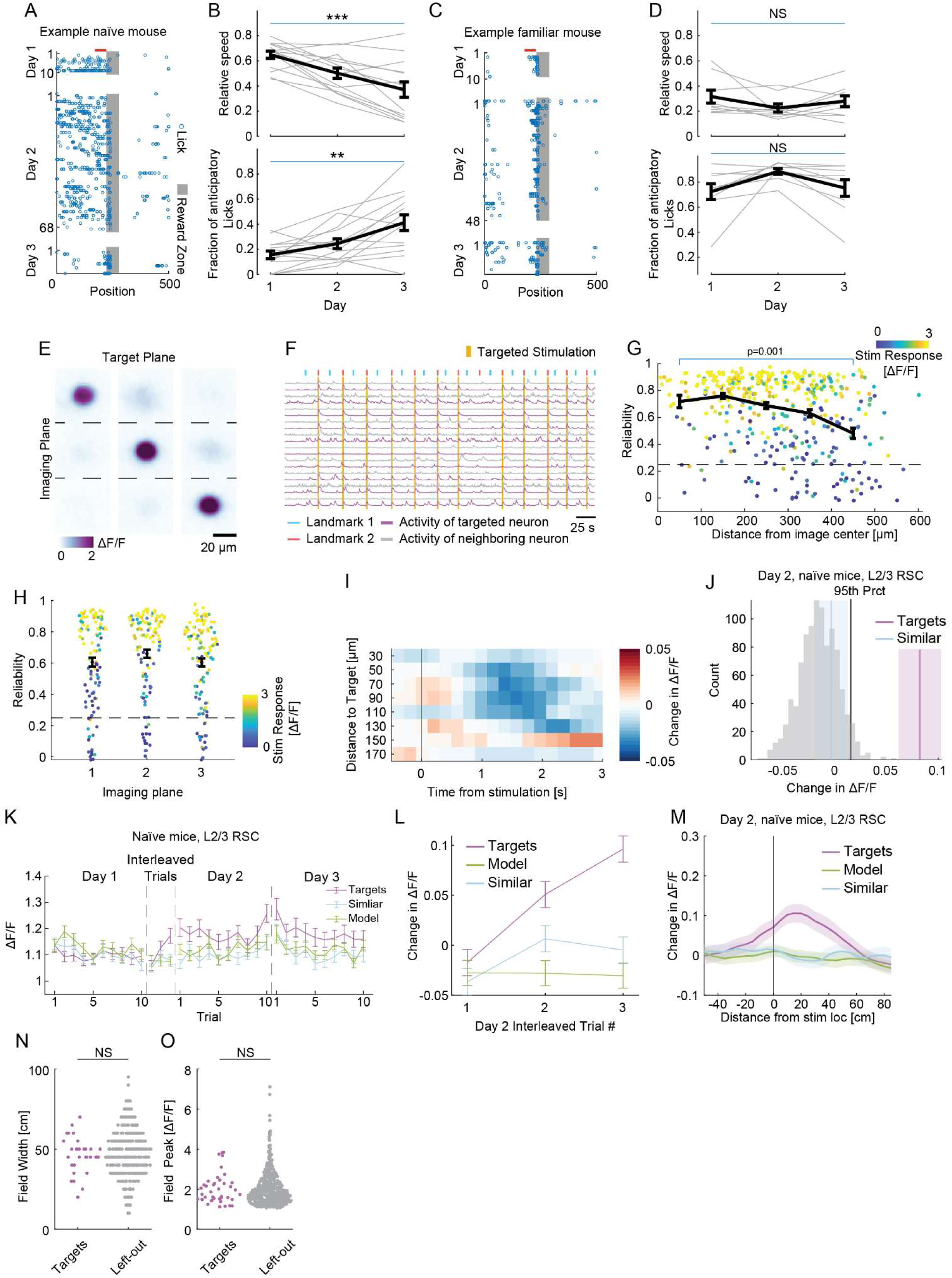
Quantification of behavior and additional quantification of stimulation evoked changes in activity, related to. **Figure 1**. A. Licking behavior of an example naïve mouse across 3 days of the experiment. Blue circles indicate individual licks, gray shading indicates the reward zone. Licks occurring immediately after the reward delivery are not shown (Methods). Red line indicates the 50 cm preceding reward zone used for quantification in (B). B. Quantification of behavior across experiment sessions in naïve mice. Top: Relative reward-approaching running speed decreased over the three days of the experiment (Methods, Day 3 vs. Day 1: p ≤ 0.0004, n = 14 mice, Wilcoxon signed-rank test). Mean ± SEM shown in black, individual animals shown in grey. Calculated as the average speed within 50 cm of the reward zone, normalized by the maximum speed over all positions preceding the reward zone. Bottom: Fraction of anticipatory licks increased over the three days, indicating improved precision in predicting reward location (Day 3 vs. Day 1: Wilcoxon signed-rank test, p ≤ 0.001, n = 14 mice). Mean ± SEM shown in black, individual animals shown in grey. Calculated as the number of non-consumatory licks within 50 cm of the reward zone (anticipatory licks) as a fraction of all non-consumatory licks (Methods). C. Same as in (A), but for the licking behavior of an example familiar mouse across 3 days of the experiment. D. Same as in (B), but for the quantification of behavior across experiment sessions in familiar mice. Top: Relative reward-approaching running speed was similar over the three days of the experiment (Day 3 vs. Day 1: p = 0.3, n = 9 mice, Wilcoxon signed-rank test). Bottom: Fraction of anticipatory licks was stable over the three days (Day 3 vs. Day 1: Wilcoxon signed-rank test, p = 0.57, n = 9 mice). Mean ± SEM shown in black, individual animals shown in grey. Calculated as the number of non-consumatory licks within 50 cm of the reward zone (anticipatory licks) as a fraction of all non-consumatory licks (Methods) E. Left column: Showing an excerpt of the average ΔF/F image of all 3 imaging planes, the excerpt is centered around all the targets located in the top imaging plane (n = 104 selected stimulation targets). Middle: Same as on the left, but for the average ΔF/F image centered around the targets located in the center imaging plane (n = 112 selected stimulation targets). Right: Same as on the left, but for targets located in the deepest imaging plane (n = 113 selected stimulation targets). Distance between imaging planes: 30 μm. F. Activity of 10 example target neurons (purple) and their closest left-out neighbor (gray) from an example naïve mouse, during stimulation trials on day 2. Orange bar indicates the targeted stimulation. Red and cyan ticks indicate the onset of landmark 1 and landmark 2. G. Stimulation reliability is constant for stimulation targets within 400 μm of the center of the field of view, and slightly decreased in stimulation targets with a distance of more than 400 μm from the center of the field of view. Color indicates the stimulation-evoked response (as in G). Dashed line indicates the threshold used to define reliably stimulated targets (Reliability > 0.3, Methods; Kruskal-Wallis test H(4) = 34.9, p ≤ 4.9*10-7, n = 329 selected stimulation targets, 5 distance bins). Stimulation targets with distance > 400 μm have significantly lower reliability (Dunn-Sidak post-hoc comparison, p < 0.04 or smaller for all groups). H. Stimulation reliability is similar across the different imaging planes (Kruskal-Wallis test H(2) = 4.68, p ≥ 0.09, n = 329 selected stimulation targets, 3 planes). I. Heatmap showing the stimulation-evoked change in activity (ΔF/F) in the left-out neurons for different distances to the closest stimulation target (median = 1083, range = 417-1803 neurons per distance bin, n = 6 mice). Activity at the onset of the stimulation-paired landmark is compared for stimulation trials and subsequent non-stimulation trials (Methods). J. Histogram of the distribution of shuffled differences in landmark-surrounding activity between Day 1 and Day 2, obtained by randomly sampling the left-out population (gray, Methods). Purple line indicates the average change in landmark-surrounding activity in Target neurons, blue line indicates the average change in activity-matched co-recorded neurons (’Similar’). Shading indicates SEM black line indicates 95th percentile (Prct) of shuffle distribution. K. Average landmark-surrounding activity for baseline trials on Day 1, interleaved trials on Day 2, post stimulation trials on Day 2, and trials on Day 3 for Target neurons (n = 260, 6 mice), and Similar and Model neurons. Error bars indicate SEM. L. Change in landmark-surrounding activity between interleaved trials (no stimulation) and baseline trials, for Target, Similar and Model neurons. Activity in Target neurons was not different from Model and Similar neurons in the first interleaved trial (p ≥ 0.05 for both, Wilcoxon signed-rank test), but increased on the second interleaved trial (Target vs Similar: p ≤ 6.3 * 10^-7^, Target vs Model: p ≤ 2.2 * 10^-25^) and the third interleaved trial (Target vs Similar: p ≤ 1.9 * 10^-19^, Target vs Model: p ≤ 1.9 * 10^-31^). Data averaged across 186 Target neurons from 4 mice. Error bars indicate SEM. M. Average change in activity between Day 2 and Day 1, surrounding the stimulation-paired landmark for Target neurons (n = 260 Target neurons, 6 mice), and Similar and Model neurons. Shading indicates SEM. N. Field width of Target neurons with a significant increase in activity and identifiable landmark-surrounding field (41 out of 55 Target neurons with significant increase in landmark-surrounding activity, Methods), and left-out neurons with an identifiable landmark-surrounding field (522 left-out neurons). We found no difference between field width (Mann-Whitney U test, p = 0.45). O. Same as M, but for peak in-field activity. We found no difference in peak amplitude between Target neurons and left-out neurons (Mann-Whitney U test, p = 0.75).

**Figure S2:**
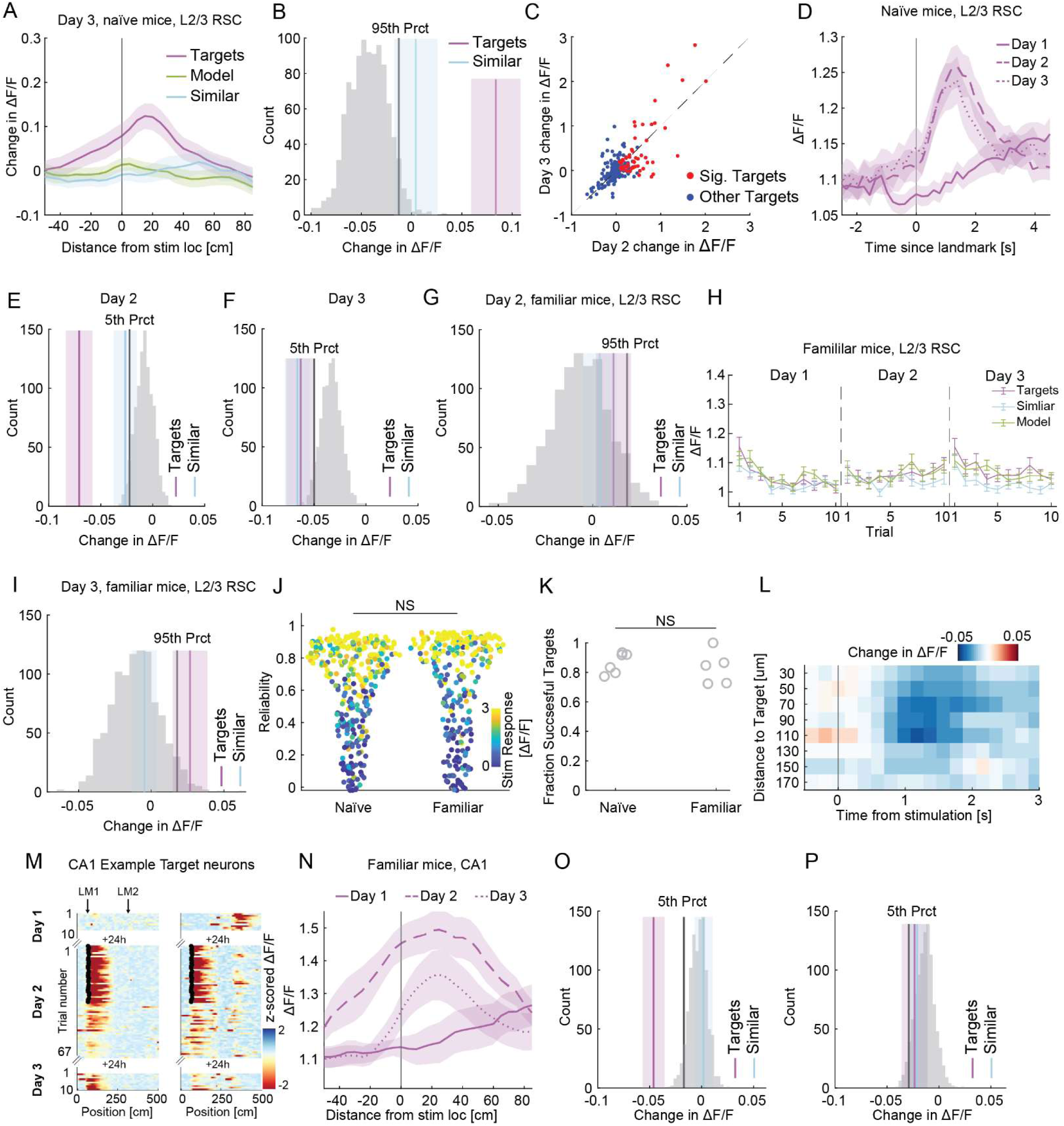
Additional quantification of changes in stimulation induced activity in RSC and CA1, related to Figure 1 and Figure 2. A. Average change in activity between Day 3 and Day 1, surrounding the stimulation-paired landmark for Target neurons (n = 260 Target neurons, 6 mice), and Similar and Model neurons. Shading indicates SEM. B. Histogram of the shuffled differences in landmark-surrounding activity between Day 1 and Day 3. Vertical lines indicate the 95th percentile of the distribution (black), and the average change in landmark-surrounding activity of Target (purple) and Similar (cyan) neurons (n = 260 Target and Similar neurons). Shading indicates SEM. C. Scatterplot showing the difference in landmark-surrounding activity on Day 2 and Day 3. Red points: Target neurons with a significant (Sig.) increase in activity on Day 2. Blue points: other Target neurons. Gray line: Identity line. Correlation between Day 2 and Day 3 = 0.73, p < 0.001, n = 260 Target neurons. D. Mean activity as a function of time around the onset of the stimulation-paired landmark of all Target neurons in naïve mice. Note that spatially binned activity is used for visualization in Fig. 1H and for quantifications throughout. Data from LM1 and LM2 Target neurons are combined (n = 260 Target neurons, 6 mice, median = 46, range = 30-56 Target neurons per mouse). Day 1 (solid line), Day 2 after stimulation trials ended (long dash) and Day 3 (short dash). Shading indicates SEM. E. Comparison of activity changes outside of the stimulation-paired landmark in Target, Similar and left-out neurons. Bars represent the average change in activity for Target neurons (purple) and Similar neurons (cyan) between post-stimulation trials on Day 2 and baseline trials on Day 1. Shading indicates SEM, obtained by averaging across all spatial positions, except for a region (- 10 cm to 60 cm) surrounding the stimulation-paired landmark. The gray histogram shows the distribution of shuffled differences for left-out neurons. The black line indicates the 5th percentile of this distribution. Activity in Target neurons is decreased more than in Similar neurons (Wilcoxon signed-rank test, p ≤ 0.001, n = 260 Target neurons). F. Same as in (E), but for the difference in activity between Day 3 and Day 1. Change in activity outside the stimulation-paired landmark in Target neurons is not different from Similar neurons (Wilcoxon signed-rank test, p ≥ 0.85). G. Histogram of the shuffled differences in landmark-surrounding activity between Day 1 and Day 2, as in (B), but for left-out neurons in RSC of familiar mice. Vertical lines indicate the 95th percentile of the distribution (black), and the average change in landmark-surrounding activity of Target (purple) and Similar (cyan) neurons (n = 208 Target and Similar neurons). H. Average landmark-surrounding activity for baseline trials on Day 1, post stimulation trials on Day 2, and trials on Day 3 for Target neurons (n = 208), and Similar and Model neurons. Error bars indicate SEM. I. Same as in (G), but for the difference in landmark-surrounding activity between Day 1 and Day 3 for left-out neurons in RSC of familiar mice. J. Scatterplot showing the distribution of stimulation reliability of all stimulation targets in naïve and familiar mice (n = 329 stimulation targets in naïve mice, n = 283 stimulation targets in familiar mice). Dots are colored by their stimulation-evoked response. No difference in average reliability and average response amplitude between naïve and familiar mice (Mann-Whitney U test, p > 0.05 for both, n = 6 naïve mice, n = 5 familiar mice). K. Scatterplot showing the fraction of stimulation targets classified as Target neurons. No difference in the fraction of successfully targeted and tracked stimulation targets between naïve and familiar mice (Mann-Whitney U test, p > 0.05, n = 6 naïve mice, n = 5 familiar mice). Gray circles: Fraction of Target neurons out of all stimulation targets per mouse. L. Heatmap showing the stimulation-evoked change in activity (ΔF/F) in the left-out neurons for different distances to the closest stimulation target (median = 1205, range = 520-1856 neurons per distance bin, n = 5 mice). Activity at the onset of the stimulation-paired landmark is compared for stimulation trials and subsequent non-stimulation trials (Methods). M. Spatially binned activity of two co-recorded Target neurons from an example CA1 mouse. Arrows on top indicate the position of stimulation-paired landmarks. Small black x’s mark the onset of stimulation. Note that activity around LM1 is significantly increased in both Target neurons (i, ii: p < 0.025, one-tailed Wilcoxon signed-rank test, comparing LM1 surrounding activity on 10 post stimulation trials to activity on Day 1). N. Mean spatially binned activity around the stimulation-paired landmark of all Target neurons in CA1 of familiar mice. Data from LM1 and LM2 Target neurons are combined (n = 78 neurons, 4 mice, median = 20, range = 16 to 22 Target neurons per mouse). Day 1 (solid line), Day 2 after stimulation trials ended (long dash) and Day 3 (short dash). Shading indicates SEM. O. Same as in (E), but for Target, Similar and left-out neurons in familiar mice. Activity outside the stimulation-paired landmarks is significantly decreased in Target neurons relative to Similar neurons (p ≤ 2.7*10^-5^, Wilcoxon signed-rank test, n = 208 neurons). P. Same as in (O), but for the difference between Day 3 and Day 1. No significant difference between Target and Similar neurons (p ≥ 0.27, Wilcoxon signed-rank test, n = 208 neurons).

**Figure S3:**
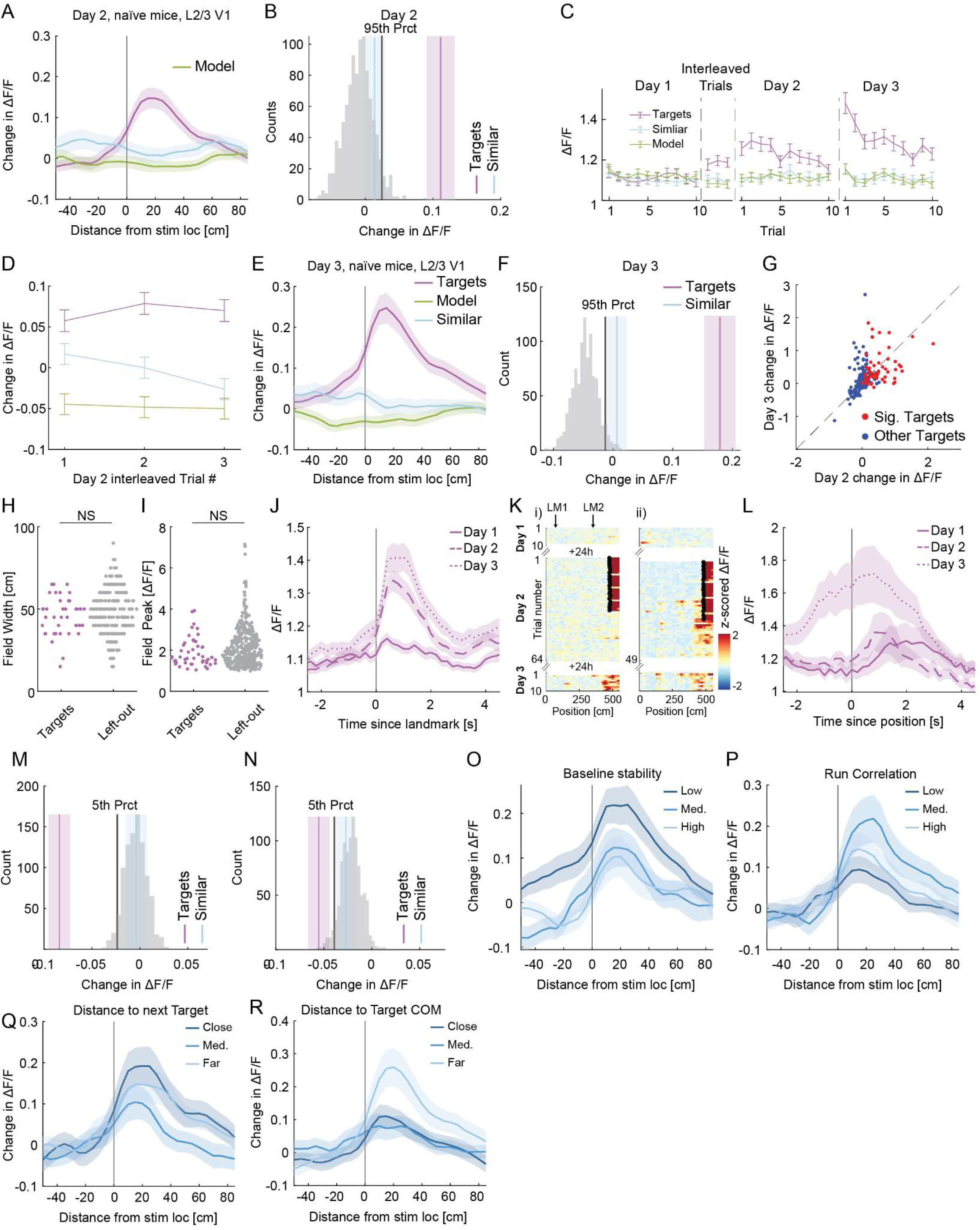
Additional quantification of stimulation evoked changes in activity in V1 of naïve mice, related to Figure 3. A. Average change in activity between Day 2 and Day 1, surrounding the stimulation-paired landmark for Target neurons in V1 of naïve mice (n = 250 Target neurons), and Similar and Model neurons. Shading indicates SEM. B. Histogram of the shuffled differences in landmark-surrounding activity between Day 1 and Day 2, but for left-out neurons in V1 of naïve mice. Vertical lines indicate the 95th percentile of the distribution (black), and the average change in landmark-surrounding activity of Target (purple) and Similar (cyan) neurons (n = 250 Target and Similar neurons). Shading indicates SEM. C. Average landmark-surrounding activity for baseline trials on Day 1, interleaved trials on Day 2, post stimulation trials on Day 2, and trials on Day 3 for Target neurons (n = 250 Target neurons), and Similar and Model neurons. Error bars indicate SEM. D. Change in landmark-surrounding activity between interleaved trials on Day 2 (no stimulation) and baseline trials, for Target, Similar and Model neurons. Activity in Target neurons is increased relative to Similar and Model neurons in all interleaved trials (Target vs Similar and Target vs Model: p ≤ 1.9 * 10^-8^ for all interleaved trials, Wilcoxon signed-rank test, n = 250 Target neurons from 8 mice). Error bars indicate SEM. E. Same as in (A), but for the difference between Day 3 and Day 1. F. Same as in (B), but for the difference between Day 3 and Day 1. G. Scatterplot showing the difference in landmark-surrounding activity on Day 2 and Day 3. Red points: Target neurons with a significant (Sig.) increase on Day 2. Blue points: other Target neurons. Gray line: Identity line. Correlation between Day 2 and Day 3: rho = 0.47, p < 0.001, n = 250 Target neurons. H. Scatterplot showing the distribution of the field width of Target neurons with significant increase in activity and identifiable landmark-surrounding field (48 out of 56 Target neurons with significant increase in landmark-surrounding activity, Methods), and left-out neurons with an identifiable field (366 left-out neurons). No difference between field width (Mann-Whitney U test, p = 0.26). I. Same as (H), but for peak in-field activity. No difference between Target neurons and left-out neurons (Mann-Whitney U test, p = 0.06). J. Mean activity as a function of time since the onset of the stimulation-paired landmark of all Target neurons in naïve mice. Data from LM1 and LM2 target neurons are combined (250 neurons, 8 mice, median = 31, range = 15 to 43 Target neurons per mouse). Day 1 (solid line), Day 2 after stimulation trials ended (long dash) and Day 3 (short dash). Shading indicates SEM. K. Spatially binned activity of two Target neurons from group 3, with stimulation paired to a position in the VR not containing strong visual landmarks (Methods). Arrows on top indicate the position of stimulation-paired landmarks for group 1 and group 2. Small black x’s mark the onset of stimulation. Note that activity around stimulation-paired position is significantly increased in both Target neurons (i and ii) on Day 3 (i, ii: p < 0.025, one-tailed Wilcoxon signed-rank test, comparing activity on 10 post stimulation trials to activity on Day 1). L. Mean activity (not spatially binned) as a function of time around the stimulation-paired position of group 3 Target neurons in a subset of naïve mice (n = 51 Target neurons, 4 mice, range = 4-20 Target neurons per mouse). Day 1 (solid line), Day 2 after stimulation trials ended (long dash) and Day 3 (short dash). Shading indicates SEM. M. Same as in (Fig. S2E), but for the difference in activity outside the stimulation-paired landmarks in L2/3 Target neurons in V1 of naïve mice. Activity in Target neurons is significantly decreased relative to Similar neurons (p ≤ 6.3 * 10^-10^, Wilcoxon signed-rank test, n = 250 Target neurons). N. Same as in (M), but for the difference between Day 3 and Day 1. No significant difference between Target and Similar neurons (p ≥ 0.07, Wilcoxon signed-rank test, n = 250 neurons). O. Average spatially binned difference between Day 2 and Day 1 in neural activity around stimulation-paired landmark for Target neurons, partitioned into 3 groups based on spatial stability during baseline trials (group 1, Low: mean stability = 0.05, n = 83 Targets, group 2, Medium: mean = 0.38, n = 82 Targets, group 3, High: mean = 0.67, n = 85 Targets). P. Same as in (O), but partitioned into 3 groups based on the correlation of neural activity with running speed during baseline trials (group 1, Low: mean correlation = -0.11, n = 83 Targets, group 2, Intermediate: mean = 0.00, n = 82 Targets, group 3, high: mean = 0.13, n = 85 Targets). Spatial stability and running speed cannot predict change in landmark-surrounding activity in Target neurons (multiple linear regression, R-squared = 0.015, F(2,245) = 1.85, p = 0.15). Q. Same as in (O), but partitioned by the planar distance to the next closest simultaneously stimulated Target neuron (group 1, close: mean distance = 33 μm, n = 83 Targets, group 2, intermediate: mean = 75 μm, n = 82 Targets, group 3, far: mean = 139 μm, n = 85 Targets). R. Same as in (O), but partitioned by mean distance to all simultaneously stimulated Target neurons (group 1, close: mean distance = 332 μm, n = 83 Targets, group 2, intermediate: mean = 409 μm, n = 82 Targets, group 3, far: mean = 502 μm, n = 85 Targets). There was no linear relationship between the change in landmark-surrounding activity and the distance to the closest Target or mean distance to all other Targets (multiple linear regression, R-squared = 0.01, F(2,247) = 1.12, p = 0.33).

**Figure S4:**
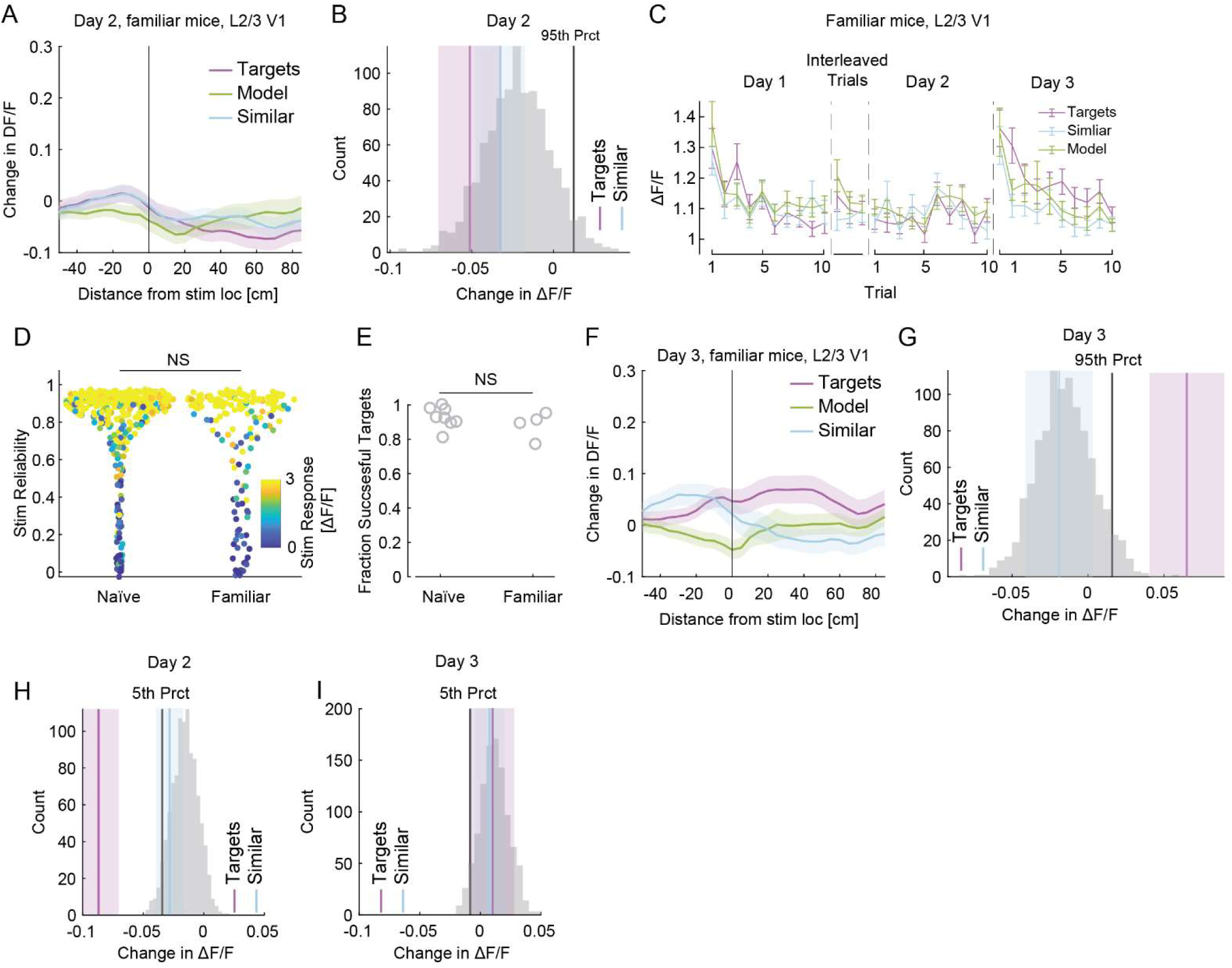
Additional quantification of stimulation evoked changes in activity in V1 of familiar mice, related to Figure 3. A. Average change in activity between Day 2 and Day 1, surrounding the stimulation-paired landmark for Target neurons in V1 of naïve mice (n = 115 Target neurons, n = 4 mice), and Similar and Model neurons. Shading indicates SEM. B. Histogram of the shuffled differences in landmark-surrounding activity between Day 1 and Day 2 for left-out neurons in V1 of familiar mice. Vertical lines indicate the 95th percentile of the distribution (black), and the average change in landmark-surrounding activity of Target (purple) and Similar (cyan) neurons. No difference in activity between Target neurons and Similar neurons (p ≥ 0.6, Wilcoxon signed-rank test, n = 115 Target and Similar neurons). Shading indicates SEM. C. Average landmark-surrounding activity for baseline trials on Day 1, interleaved trials on Day 2, post stimulation trials on Day 2, and trials on Day 3 for Target neurons (n = 115 neurons, 4 mice), and Similar and Model neurons. Error bars indicate SEM. D. Scatterplot showing the distribution of stimulation reliability of all stimulation targets in naïve and familiar mice (n = 338 stimulation targets in naïve mice, n = 174 stimulation targets in familiar mice). Individual dots are colored by the stimulation-evoked response in each stimulation target. No difference in average reliability and average response amplitude between naïve and familiar mice (Mann-Whitney U test, p > 0.05 for both, n = 7 naïve mice, n = 4 familiar mice). E. No difference in the fraction of successfully targeted and tracked stimulation targets between naïve and familiar mice (Mann-Whitney U test, p > 0.05, n = 7 naïve mice, n = 4 familiar mice). Gray circles: Fraction of Target neurons out of all stimulation targets per mouse. F. Same as in (A), but for the change in activity between Day 3 and Day 1. G. Same as in (B), but for the change in activity between Day 3 and Day 1. Difference in activity is significantly larger in Target neurons compared to Similar neurons (p ≤ 3.1*10^-5^, Wilcoxon signed-rank test, n = 115 Target and Similar neurons). H. Same as in (Fig. S2E), but for the difference in activity outside the stimulation-paired landmarks in L2/3 Target neurons in V1 of familiar mice. Activity in Target neurons is significantly decreased relative to Similar neurons (p ≤ 0.0009, Wilcoxon signed-rank test, n = 115 Target neurons) I. Same as in (H), but for the difference between Day 3 and Day 1. No significant difference between Target and Similar neurons (p ≥ 0.55, Wilcoxon signed-rank test, n = 115 Target neurons).

**Figure S5:**
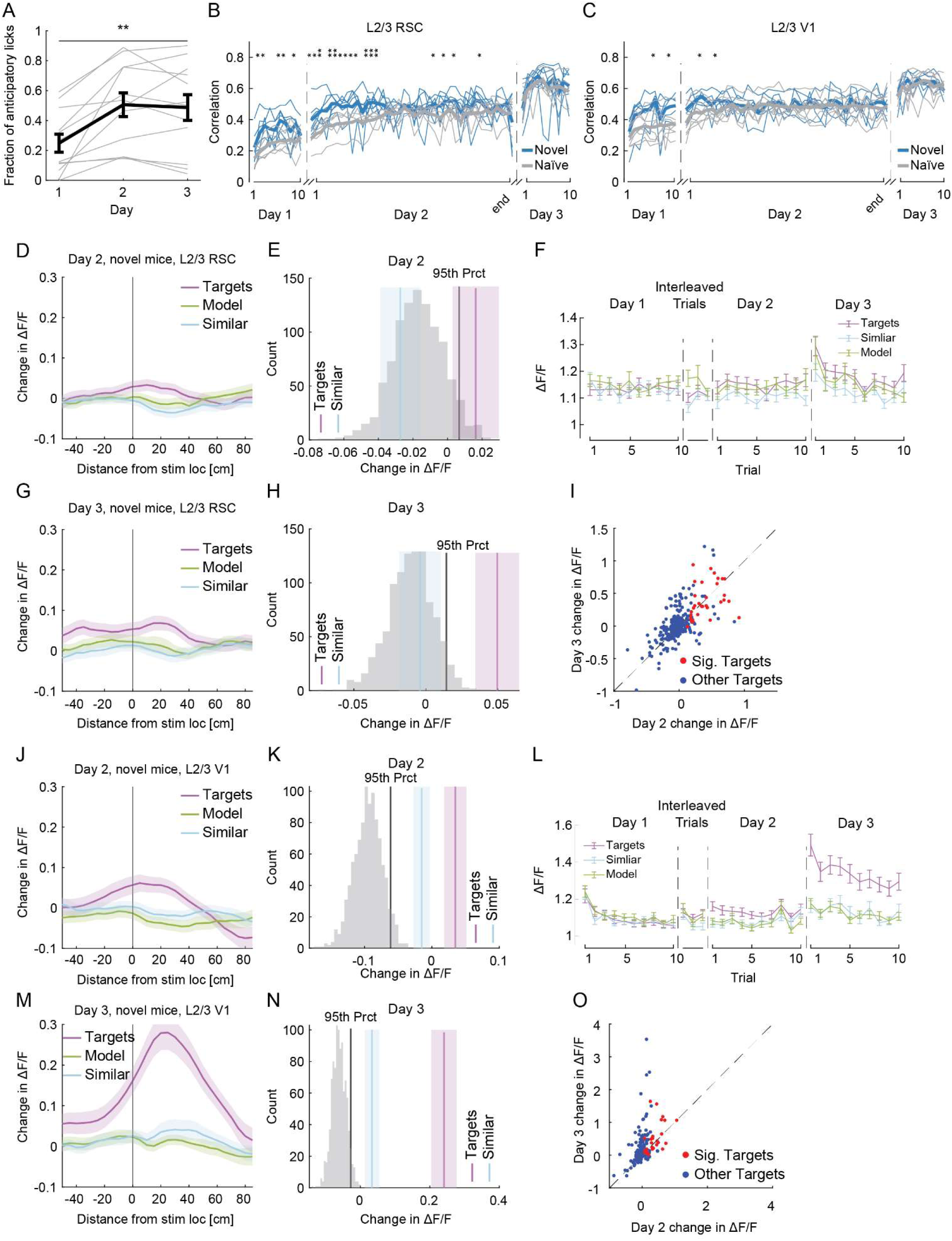
Additional quantification of behavior and neural activity of novel mice. Related to Figure 4. A. The fraction of anticipatory licks increased over the three days, when switching from a familiar to a novel environment (Day 3 vs. Day 1: Wilcoxon signed-rank test, p ≤ 0.009, n = 12 novel mice). Mean ± SEM shown in black, individual animals shown in grey. B. Correlation of spatial activity patterns of left-out neurons in RSC between Days 1 and 2 and Day 3 in naïve (gray) and novel (blue) mice. Higher correlation indicates greater similarity to Day 3. Thick lines: mean; thin lines: individual mice. *p < 0.05, **p < 0.01 (Mann-Whitney U test, n = 6 naïve mice, 7 novel mice). Day 2 data is interpolated to accommodate varying trial numbers (Methods). C. Same as in (B), but comparing correlation of spatial activity patterns between left-out neurons in V1 of naïve and novel mice (n = 7 naïve mice, n = 5 novel mice). D. Average change in activity between Day 2 and Day 1, surrounding the stimulation-paired landmark for Target neurons, Similar and Model neurons in RSC of novel mice (n = 303 Target neurons). Shading indicates SEM. E. Histogram of the shuffled differences in landmark-surrounding activity between Day 1 and Day 2 for left-out neurons in RSC of novel mice. Vertical lines indicate the 95th percentile of the distribution (black), and the average change in landmark-surrounding activity of Target (purple) and Similar (cyan) neurons (n = 303 Target and Similar neurons). Shading indicates SEM. F. Average landmark-surrounding activity for baseline trials on Day 1, interleaved trials on Day 2, post stimulation trials on Day 2, and trials on Day 3 for Target neurons (n = 303 Target neurons), and Similar and Model neurons. Error bars indicate SEM. G. As in (D), but for the difference between Day 3 and Day 1. H. As in (E), but for the difference between Day 3 and Day 1. I. Scatterplot showing the difference in landmark-surrounding activity on Day 2 and Day 3. Red points: Target neurons with a significant increase on Day 2. Blue points: other Target neurons. Gray line: Identity line. Correlation between Day 2 and Day 3 = 0.67, p < 0.001, n = 303 Target neurons. J.-N. as in D-H, but for Target neurons in V1 of novel mice (n = 192 Target neurons, 5 novel mice). O. Scatterplot showing the difference in landmark-surrounding activity on Day 2 and Day 3 in Target neurons. Red points: Target neurons with a significant increase on Day 2. Blue points: other Target neurons. Gray line: Identity line. Correlation between Day 2 and Day 3: rho = 0.43, p < 0.001, n = 192 Target neurons. Landmark-surrounding activity is significantly larger on Day 3, compared to Day 2 (Wilcoxon signed-rank test, p ≤ 0.01, n = 192 Target neurons).

**Figure S6:**
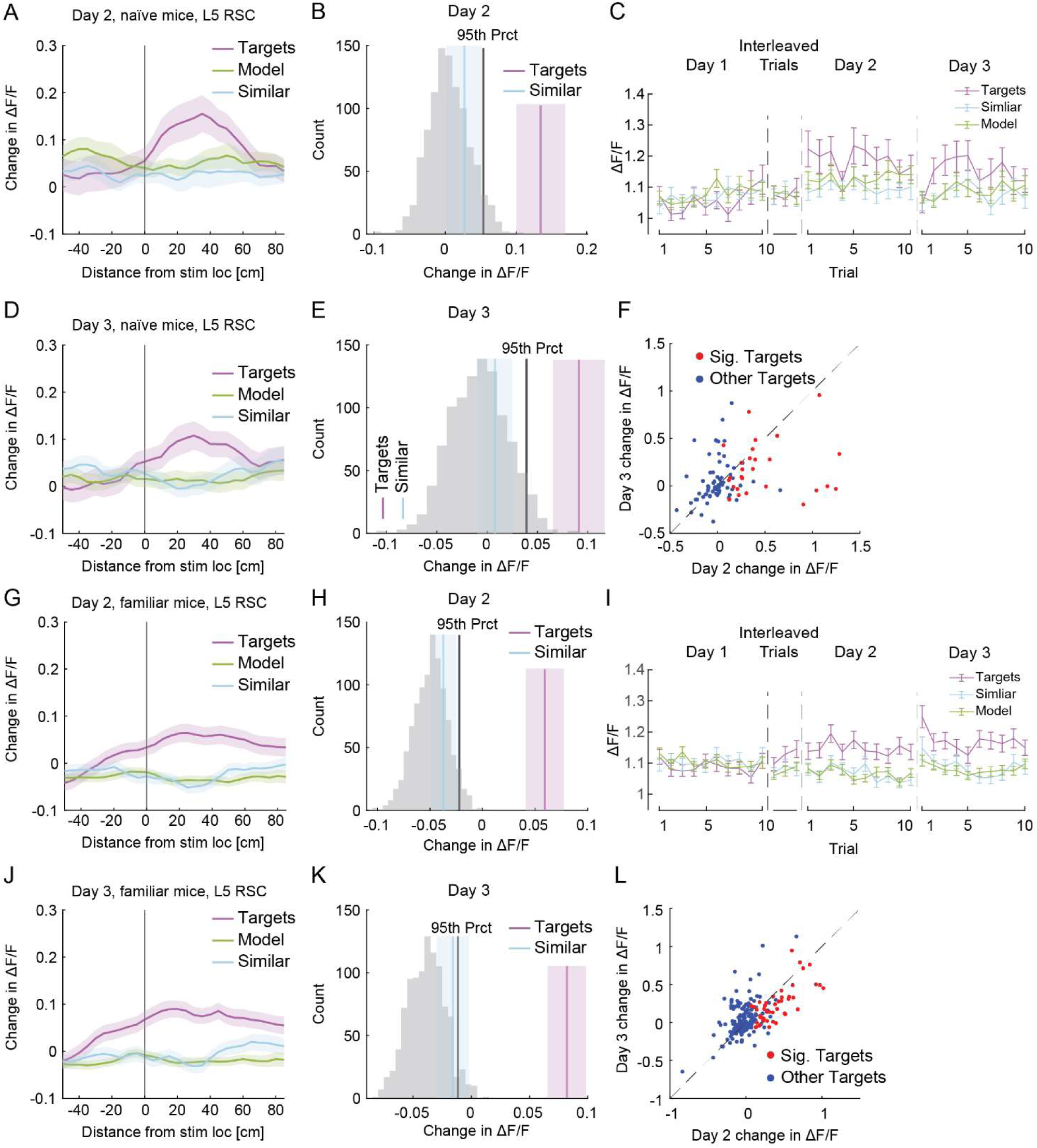
Additional quantification of stimulation evoked changes in activity of Layer 5 Target neurons from RSC of naïve and familiar mice. A. Average change in activity between Day 2 and Day 1, surrounding the stimulation-paired landmark for layer 5 (L5) Target neurons (n = 94 Target neurons), and Similar and Model neurons in RSC of naïve mice. Shading indicates SEM. B. Histogram of the shuffled differences in landmark-surrounding activity between Day 1 and Day 2 for left-out L5 neurons in RSC of novel mice. Vertical lines indicate the 95th percentile of the distribution (black), and the average change in landmark-surrounding activity of Target (purple) and Similar (cyan) neurons. Difference in activity is significantly larger in Target neurons, compared to Similar neurons (p ≤ 0.01, Wilcoxon signed-rank test, n = 94 Target and Similar neurons). Shading indicates SEM. C. Average landmark-surrounding activity for baseline trials on Day 1, interleaved (non-stimulation) trials on Day 2, post stimulation trials on Day 2, and trials on Day 3 for Target neurons (n = 94 Target neurons), and Similar and Model neurons. Error bars indicate SEM. D. As in (A), but for the difference between Day 3 and Day 1. E. As in (B), but for the difference between Day 3 and Day 1. Difference in activity is significantly larger in Target neurons, compared to Similar neurons (p ≤ 0.02, Wilcoxon signed-rank test, n = 94 Target and Similar neurons). F. Scatterplot showing the difference in landmark-surrounding activity on Day 2 and Day 3. Red points: Target neuron with a significant increase on Day 2. Blue points: other Target neurons. Gray line: Identity line. Correlation between Day 2 and Day 3: rho = 0.30, p < 0.001, n = 94 Target neurons. G.-K. Same as A-E, but for Target neurons in L5 of RSC (n = 205 Target neurons, 6 familiar mice). Difference in activity between Day 2 and Day 1 is significantly larger in Target neurons, compared to Similar neurons (p ≤ 6.2*10^-5^, Wilcoxon signed-rank test, n = 205 Target and Similar neurons), and significantly larger between Day 3 and Day 1 (p ≤ 5.5*10^-5^, Wilcoxon signed-rank test, n = 205 Target and Similar neurons). L. Scatterplot showing the difference in landmark-surrounding activity on Day 2 and Day 3 in Target neurons. Red points: Target neuron with a significant increase on Day 2. Blue points: other Target neurons. Gray line: identity line. Correlation between Day 2 and Day 3: rho = 0.62, p < 0.001, n = 205 Target neurons.

